# Innate immune stress pathway activation underlies heterochromatin dysfunction pathology

**DOI:** 10.64898/2026.09.23.753861

**Authors:** Roopali Pradhan, Anna F Townley, Anna V Protasio, Joshua Miguel Danac, Yan Dong, Yushan Fang, Alex Appert, Francesco N Carelli, Shixun Han, Heli Vaikkinen, Iva Tchasovnikarova, Julie Ahringer

## Abstract

Heterochromatin loss disrupts nuclear architecture, gene regulation and repetitive element silencing, and is associated with diverse human diseases. However, mechanisms linking heterochromatin dysfunction to pathological phenotypes remain unclear. Using genetic interaction screening and genomic analyses in *C. elegans*, we identify secondary activation of the Intracellular Pathogen Response (IPR), an innate immune stress pathway, as a major contributor to heterochromatin mutant phenotypes. Constitutive IPR activation phenocopies slow growth and indirect transcriptional changes observed in these mutants. Depletion of genetic enhancers further increased, whereas suppressor RNAi attenuated IPR activation, with direct heterochromatin targets remaining substantially deregulated. Notably, many suppressors encode active chromatin components, and mild reduction of RNA polymerase II activity ameliorates growth defects in *C. elegans* HP1 mutants and human HP1-deficient cells. Our findings reveal secondary stress response activation as an important mechanism linking heterochromatin dysfunction to pathology and identify transcriptional dampening as a potential therapeutic strategy for mitigating these effects.

## Main

Heterochromatin is a fundamental component of eukaryotic genomes that plays crucial roles in genome regulation and nuclear organization. Characterized by its condensed structure and specific histone modifications, particularly histone H3 lysine 9 methylation (H3K9me), heterochromatin comprises a significant fraction of most genomes and is essential for silencing genes and repetitive elements, maintaining genome stability, and correct nuclear architecture^1^. A hallmark of animal heterochromatin is heterochromatin protein 1 (HP1), which binds H3K9me2/3 through its chromodomain and contributes to the organization and maintenance of repressive chromatin. The loss or dysregulation of heterochromatin has been associated with numerous human diseases, including cancer, neurodegeneration, and premature aging syndromes, highlighting its importance for normal cellular function and organismal health^2,3^. However, despite its clinical relevance, the mechanisms by which heterochromatin dysfunction leads to pathological outcomes remain poorly understood. This knowledge gap presents a barrier to developing therapeutic strategies for heterochromatin-associated diseases.

*C. elegans* has two HP1 orthologs, HPL-1 and HPL-2, together with many other conserved heterochromatin factors^4^. HPL-2 functions together with the multi-zinc finger protein LIN-13 and the L3MBTL2 ortholog LIN-61, which physically and genetically interact with HPL-2, while the SETDB1 ortholog MET-2 generates H3K9me1/2 ^5,6^. HPL-2, LIN-13, LIN-61 and MET-2 show highly similar genomic distributions that closely correspond to H3K9me2, and loss of these factors causes overlapping deregulation of gene and repetitive element expression^5–10^. Mutations in these heterochromatin genes also cause a shared spectrum of phenotypes, including temperature-sensitive slow growth, fertility defects, developmental abnormalities and inappropriate expression of germline-specific genes in the soma^5,9–17^. Moreover, double-mutant combinations show enhanced phenotypes compared with single mutants, and genetic suppressors of heterochromatin mutant phenotypes have been identified^5,9,10,14–16,18–20^. Together, these observations suggest that conserved heterochromatin factors act in an interconnected functional network, and that their loss may produce phenotypes through shared downstream mechanisms.

A major unresolved question is whether the phenotypes caused by heterochromatin loss arise primarily from direct derepression of heterochromatin targets, such as genes and repetitive elements, or from secondary cellular states triggered by this deregulation. Distinguishing these possibilities is important because indirect pathological states may be more amenable to suppression than the primary chromatin defect itself. We therefore reasoned that large-scale screening for enhancers and suppressors of heterochromatin mutant phenotypes would provide a powerful approach to identify the functional network of heterochromatin and uncover the physiological basis of heterochromatin dysfunction phenotypes.

To this end, we performed RNAi genetic interaction screens in five heterochromatin mutant backgrounds. Analyses of the enhancer and suppressor networks strongly implicate stress response activation as a major functional component of the physiological defects associated with heterochromatin loss. In particular, we found that heterochromatin mutants activate the Intracellular Pathogen Response (IPR), and that genetic activation of this pathway is sufficient to reproduce slow growth and many indirect gene expression changes observed in heterochromatin mutants. Many suppressor genes encode components of active chromatin complexes, whose inhibition preferentially attenuates IPR-associated transcriptional changes, whereas inhibition of enhancer genes amplifies these changes. Finally, mildly reducing RNA polymerase II activity improves heterochromatin mutant defects in both *C. elegans* and human HP1-deficient cells. Thus, modulation of transcriptional activity or conserved suppressor pathways may provide potential therapeutic avenues for heterochromatin-associated diseases.

## Results

### Generation of a heterochromatin genetic interaction network

We selected five heterochromatin mutants for genetic interaction screening: *hpl-1/*HP1*, hpl-2/*HP1*, lin-13, lin-61/*L3MBTL2, and *met-2/*SETDB1 (Fig. 1a). These mutants display temperature-dependent growth retardation, which varies in severity across the mutants (Extended Data Fig. 1a). To enable the throughput needed for RNAi screening of multiple mutants, we used an RNAi sub-library that targets 2292 genes encoding nuclear proteins (Fig. 1a and Supplementary Table 1). We screened all five mutants for enhancers, defined as RNAi treatments that worsened growth or fertility relative to the effect of the same RNAi treatment in wild type. Suppressor screening was limited to *hpl-2* and *lin-13*, as only their slow growth phenotypes were strong enough to enable reliable detection of rescue. RNAi was initiated in L3 larvae, and effects were assayed in those worms (P0) and their progeny (F1) (Fig. 1a).

**Figure 1.**
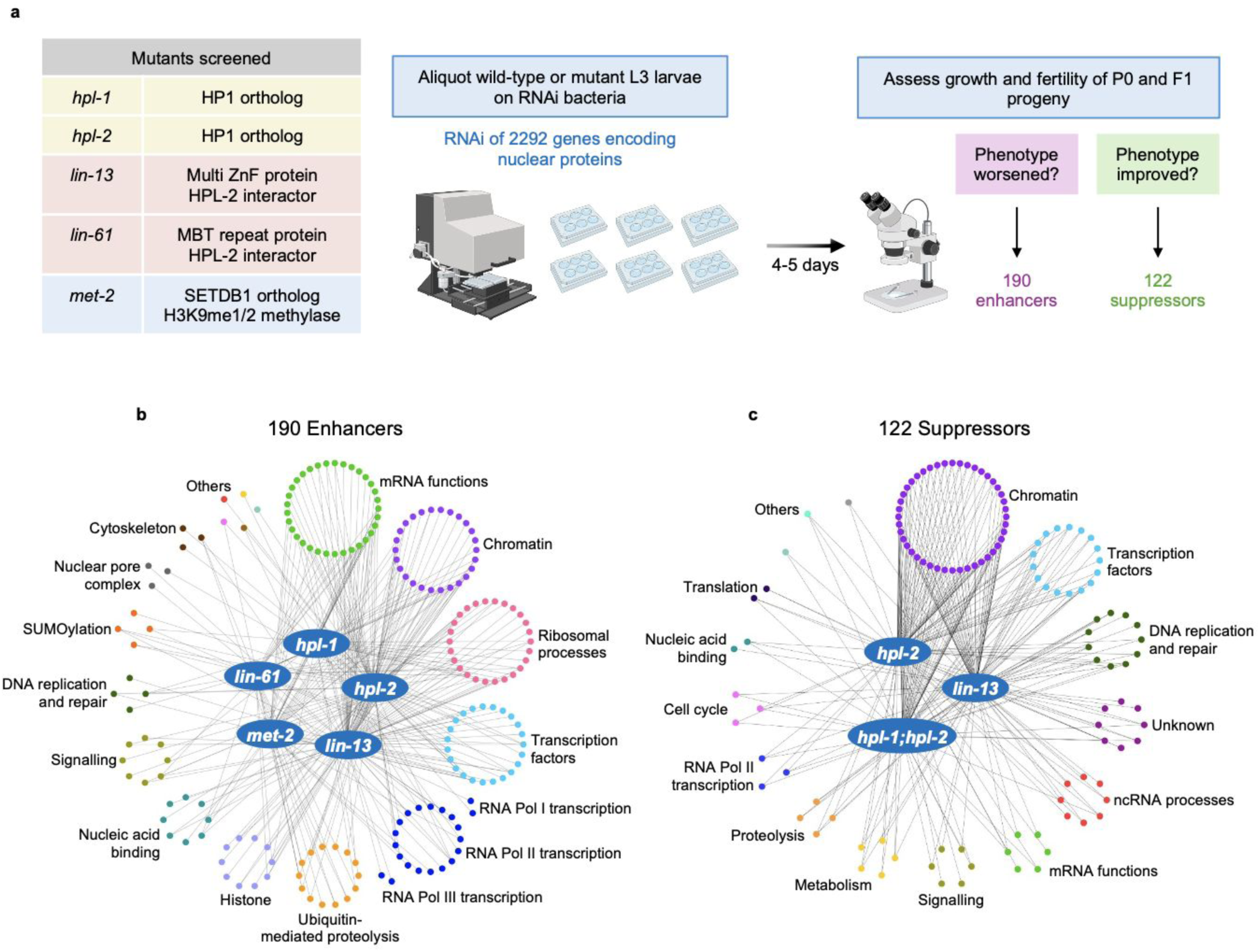
Genetic interaction screens identify enhancers and suppressors of heterochromatin mutant phenotypes. **a**, Experimental design of genetic interaction screens. RNAi of 2292 genes encoding nuclear proteins was carried out in five heterochromatin mutants (*hpl-1, hpl-2, lin-13, lin-61, and met-2*) and phenotypic assessment was performed to identify enhancers and suppressors (Figure created using Biorender.com). **b**,**c**, Genetic interaction networks of 190 enhancers (**b**) and 122 suppressors (**c**) of heterochromatin mutants (central blue target nodes). Genetic interactors are organised by functional class. Lines connect enhancers and suppressors to their mutant nodes.

Following two rounds of validations, we identified 190 genetic enhancers where RNAi knockdown worsened a heterochromatin mutant phenotype, and 122 suppressors where RNAi knockdown caused phenotypic rescue (Fig. 1b,c and Supplementary Table 2). The most common enhancement phenotype was a slowing of growth rate (71% of enhancers). The genetic interaction networks have high interconnectivity, supporting shared functions of the heterochromatin genes, with the majority of each mutant’s enhancers (59-85%) also enhancing the phenotype of at least one other mutant (Fig. 1b,c and Supplementary Table 2). Of the suppressors identified, 57% were shared between *hpl-2* and *lin-13*. Furthermore, 63% of suppressors also rescued the slow growth of *hpl-1; hpl-2* double mutants, which lack both HP1 orthologues and have the most severe growth defect among the heterochromatin mutants that we tested (Fig. 1c; Extended Data Fig. 1a and Supplementary Table 2).

The enhancer and suppressor genes have diverse predicted functions (Fig. 1b,c and Supplementary Table 2). Of note, genes involved in ribosomal processes such as ribosome biogenesis and in protein modification pathways such as SUMOylation were found among enhancers but not suppressors (Fig. 1b,c). Additionally, genes involved in ubiquitin-mediated proteolysis were more associated with enhancers, whereas genes involved in non-coding RNA processes were more associated with suppressors. Further, although chromatin functions were found among enhancer and suppressor genes, the types of functions differed. Enhancers had frequent annotations for SynMuv class genes, many of which have transcription repression functions^21^, whereas suppressor genes more often encoded proteins in chromatin complexes associated with active chromatin, such as components of the NSL, COMPASS, TIP60, and ISWI complexes (Supplementary Table 2).

Consistent with the enrichment of ribosome biogenesis genes among enhancers, the heterochromatin mutants displayed smaller nucleoli and reduced rRNA content, together with increased eIF2α phosphorylation, indicating activation of the Integrated Stress Response (ISR) pathway, which slows down translation in response to proteotoxic stress^22,23^ (Extendedended Data Figs. 1c-f and 2a,b). ISR activation appears to be protective rather than causal for these phenotypes, because preventing eIF2α phosphorylation did not restore rRNA levels and exacerbated the growth defect of *hpl-2* mutants (Extended Data Fig. 2c,d).

**Figure 2.**
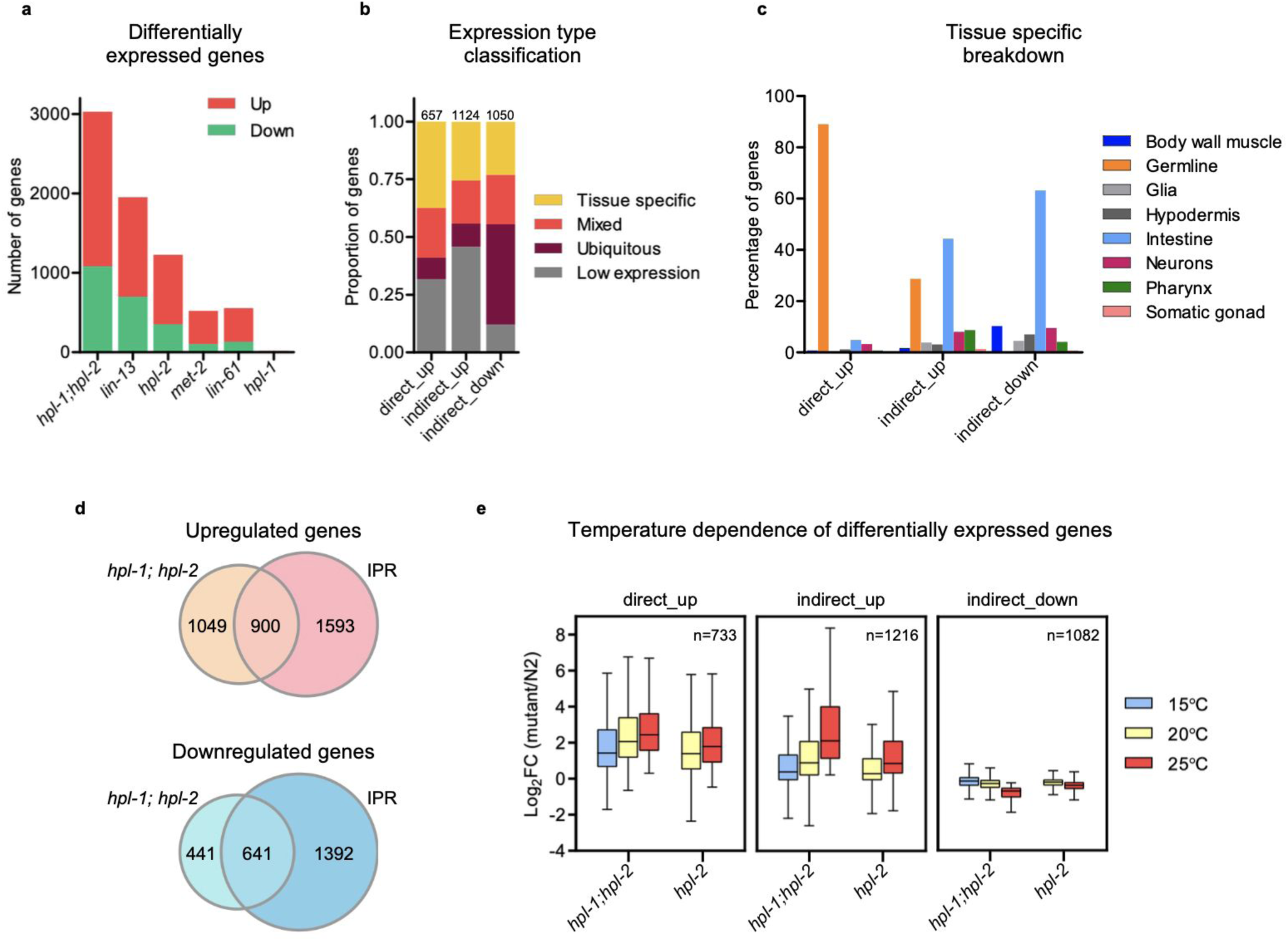
Direct and indirect transcriptional effects of heterochromatin loss. **a**, Number of up- and downregulated genes in each heterochromatin mutant (adjusted p-value: *p* <0.001). **b**, Classification of *hpl-1; hpl-2* direct_up, indirect_up and indirect_down genes into different expression types (tissue specific, mixed, ubiquitous or low expression; see methods for definitions). **c**, Tissue of maximum expression for tissue-specific *hpl-1; hpl-2* direct_up, indirect_up and indirect_down genes from (**b**). **d**, Venn diagram showing the overlap between *hpl-1; hpl-2* upregulated genes and the IPR_up signature (top), and *hpl-1; hpl-2* downregulated genes and the IPR_down signature (bottom). **e**, Box plots showing the extent of deregulation of *hpl-1; hpl-2* and *hpl-2* direct_up, indirect_up and indirect_down genes at different temperatures.

### Heterochromatin loss causes extensive direct and indirect transcriptional changes

To investigate potential causes of heterochromatin mutant defects, we assessed and compared the nature of gene expression deregulation in the mutants. We performed RNA-seq on starved L1 larvae to match stages across samples, as the mutants have different growth rates and also often grow asynchronously (Extended Data Fig. 1a). Assessing L1 larvae also enabled detection of germline gene expression in the soma, which was previously reported for several SynMuv B mutants, including *hpl-2, lin-13, lin-15B,* and *lin-35*^5,16,24–26^. L1s contain only two germline precursor cells among their 558 cells and so are essentially comprised of somatic cells. We profiled all single mutants included in the RNAi screens as well as *hpl-1; hpl-2* double mutants.

In line with a loss of repressive heterochromatin, we detected more upregulated than downregulated genes across the mutant strains (Fig. 2a and Supplementary Table 3). An exception was *hpl-1*, for which nearly no gene expression changes were detected, consistent with its predominantly normal growth and fertility^27^; these data were not analysed further. Notably, supporting the previously reported redundancy between *hpl-1* and *hpl-2* ^12^, we observed more extensive gene expression changes in *hpl-1; hpl-2* double mutants compared to the single mutants (Fig. 2a).

The other four heterochromatin mutant strains (*hpl-2, lin-13, met-2, lin-61*) had highly similar gene expression alterations, and genes up- or down-regulated in the single mutants are largely subsets of the deregulated genes in *hpl-1; hpl-2* mutants (Extended Data Fig. 3a and Supplementary Table 3). We also found that the slower growing strains (*hpl-1; hpl-2*, *hpl-2*, and *lin-13*) have more gene expression changes compared to those with mild growth defects (*lin-61* and *met-2*) (Fig. 2a and Extended Data Fig. 1a), suggesting a possible relationship between gene expression changes and growth rate. We focused further analyses on *hpl-1; hpl-2* double mutants as the gene expression changes in that strain are the most extensive and encompass the vast majority of changes seen across all single mutants (Extended Data Fig. 3a).

We considered that transcriptional changes caused by loss of heterochromatin function would result from a mixture of direct loss of repression at heterochromatin-marked genes and indirect effects of heterochromatin loss, and that comparing these classes should be informative. To identify heterochromatic genomic regions, we performed ChIP-seq of HPL-2, LIN-13, LIN-61, MET-2, H3K9me2 and H3K9me3 in starved L1 larvae. As previously observed in adults^10^, the binding patterns of the different heterochromatin proteins were highly similar to each other and to the pattern of H3K9me2 (Extended Data Fig. 4a,b and Supplementary Table 4). Given the concordance of H3K9me2 with heterochromatin factor binding and variations in the signal to noise among the heterochromatin ChIP-seq datasets, likely due to differences in antibody quality, we used promoter H3K9me2 as an operational criterion to distinguish direct targets of heterochromatin repression from genes whose altered expression arises indirectly. Among genes upregulated in *hpl-1; hpl-2* mutants, those with H3K9me2-marked promoters were classified as direct targets and termed “direct_up” (n=733), whereas those lacking promoter H3K9me2 were classified as indirect upregulated targets, “indirect_up” (n=1216). Since the loss of a repressive chromatin pathway would not be expected to directly reduce transcription, downregulated genes were classified as “‘indirect_down” (n=1082) (Supplementary Table 5).

We first investigated the tissue specificity of expression of direct_up, indirect_up and indirect_down genes by separating each group into four expression type classes: tissue-specific, mixed, ubiquitous and low expression (Fig. 2b and Supplementary Table 6; see methods). We found that the tissue-specific indirect_up and indirect_down genes were most biased for intestinal expression (Fig. 2c), suggesting that a major indirect consequence of heterochromatin loss is a change in intestinal gene expression. Interestingly, nearly all of the tissue-specific direct_up genes and a large fraction of the indirect_up genes have germline-biased expression (Fig. 2c). As the profiled starved L1 larvae have essentially no germline tissue, this indicates that germline genes become expressed in somatic tissues, as has been previously observed for *hpl-2* mutants and in mutants of other chromatin repressors^5,16,17,24–26^. The presence of germline-biased genes in both the direct_up and indirect_up classes suggests that ectopic germline gene expression in the soma arises through both direct loss of heterochromatin-mediated repression and secondary transcriptional changes.

### Heterochromatin loss indirectly induces the Intracellular Pathogen Response (IPR) stress pathway

To further investigate the nature of the direct and indirect transcriptional changes, we performed functional enrichment analyses using WormCat^28^. Direct_up genes were enriched in small RNA pathways, pseudogenes, nucleic acid binding, and proteins containing both MATH and BTB domains, a class of substrate adaptors for Cullin-RING E3 ubiquitin ligases^29^ (Extended Data Fig. 3b). Both direct_up and indirect_up genes were enriched for functions related to proteolysis and chromosome dynamics, whereas indirect_down genes were enriched for ribosome subunits and diverse metabolic functions, including amino acid, glycolytic, mitochondrial, lipid and nucleotide metabolism (Extended Data Fig. 3b). These latter enrichments, together with the reduced rRNA content of *hpl-1; hpl-2* mutants, suggest broad reductions in translational and metabolic activity. Most notably, the strongest enrichment among indirect_up genes was for proteins containing an ALS2cr12 domain.

The ALS2cr12 domain is a defining feature of the *pals* gene family^30,31^. A set of 80 highly induced genes, including 22 *pals* genes, is widely used as an operational signature of activation of the Intracellular Pathogen Response (IPR), an innate immune stress pathway induced by diverse triggers such as intracellular pathogens, viral infection, and proteotoxic stress^32–34^. The IPR has features in common with the mammalian type I interferon response, including sensing of viral RNA through a RIG-I-like receptor and outcomes of pathogen defence and impaired development^35–37^. The 80-gene signature provides a marker of IPR activation, but the broader transcriptional consequences of IPR pathway activation have not been defined.

Constitutive IPR activation can itself impair development. *pals-17* and *pals-22* encode negative regulators of the IPR, and loss of either causes upregulation of the 80-gene IPR signature and growth defects, with *pals-17* showing stronger effects^33,38^. We therefore considered whether IPR activation might contribute to the growth defects caused by heterochromatin loss. We found that *pals-17* mutants and *hpl-1; hpl-2* double mutants had similarly strong temperature-dependent slow growth defects, whereas *pals-22* mutants had weaker growth defects (Extended Data Fig. 1b). *pals-17* mutants also had small nucleoli, reduced rRNA levels and weak ISR activation, phenotypes shared with heterochromatin mutants (Extended Data Figs. 1d-f and 2a,b).

To define the broader transcriptional consequences of IPR activation, we profiled gene expression in starved L1 *pals-17* mutants, matching the developmental stage used for heterochromatin mutant profiling. A previous RNA-seq analysis of *pals-17* was performed at later larval stages, but developmental asynchrony meant it was not possible to distinguish IPR-associated transcriptional changes from differences in developmental stage^38^. To assess whether the changes observed in *pals-17* reflected IPR pathway activation rather than *pals-17*-specific effects, we also profiled *pals-22* mutants. *pals-22* showed a weaker but concordant transcriptional response: most genes significantly deregulated in *pals-22* were also deregulated in *pals-17*, and genes reaching significance in only one mutant showed concordant changes as a group in the other mutant (Extended Data Fig. 5a-c). As *pals-17* produced the stronger transcriptional response and, like *hpl-1; hpl-2*, a strong temperature-sensitive growth defect, we used genes deregulated in *pals-17* to define the broader IPR-associated transcriptional state for subsequent analyses. We termed genes significantly up- or downregulated in *pals-17* mutants IPR_up (n = 2,493) or IPR_down (n = 2,033), respectively (Supplementary Table 3).

Strikingly, the IPR_up and IPR_down signatures showed extensive overlap with transcriptional changes in heterochromatin mutants (Fig. 2d and Extended Data Fig. 3c); 46% of genes upregulated and 59% of genes downregulated in *hpl-1; hpl-2* were also deregulated upon IPR activation. Moreover, 84% of these shared genes fell with the indirect classes (Extended Data Fig. 5d). IPR_up genes predominantly showed germline- or intestinal-biased expression, whereas IPR_down genes were primarily intestinally enriched, closely mirroring the *hpl-1; hpl-2* indirect_up and indirect_down classes, respectively (Fig. 2c and Extended Data Fig. 5e,f). The two sets also shared functional enrichments (Extended Data Fig. 3b), with IPR_down genes accounting for the majority of metabolic downregulation observed in *hpl-1; hpl-2* mutants. Thus, constitutive IPR activation produces a broad transcriptional state that closely recapitulates the indirect response to heterochromatin loss, including downregulation of metabolic and translational functions.

We next asked whether the IPR-associated state tracks the temperature-dependent severity of heterochromatin mutant growth defects. Gene expression profiling of *hpl-2* and *hpl-1; hpl-2* mutants at different temperatures showed that IPR-associated indirect transcriptional changes preferentially increased at higher temperature, paralleling the worsening growth defects of these mutants (Fig. 2e; Extended Data Fig. 5g and Supplementary Table 7). Together, the phenotypic similarity, extensive transcriptional concordance and temperature dependence support indirect activation of the IPR as a major contributor to the growth defects caused by heterochromatin loss.

### Genetic suppressors and enhancers preferentially affect the IPR-associated state

We next asked whether altered IPR activation underlies the opposing effects of suppressor and enhancer depletion on heterochromatin mutant growth. To this end, we first analysed gene expression changes after RNAi of 14 suppressors in *hpl-1; hpl-2* double mutants. The suppressors included components of the NSL and TIP60 histone acetyltransferase complexes, the COMPASS complex, the PRC2 complex, the ISWI nucleosome remodelling complex, histone HTZ-1/H2A.Z, and the ubiquitin peptidase MATH-33/USP7. RNAi of each suppressor substantially ameliorated the transcriptional alterations in *hpl-1; hpl-2* mutants. In all cases, indirect targets were more strongly corrected than direct targets, and the 80-gene IPR signature was significantly reduced (Fig. 3a,c; Extended Data Fig. 6a and Supplementary Table 8). Notably, the suppression of indirect target gene misexpression predominantly occurred in the subset overlapping with IPR pathway activation (Fig. 3a). None of the 14 suppressor RNAi treatments rescued the growth defect of *pals-17* mutants (Supplementary Table 9), indicating that they act upstream of IPR activation.

**Figure 3.**
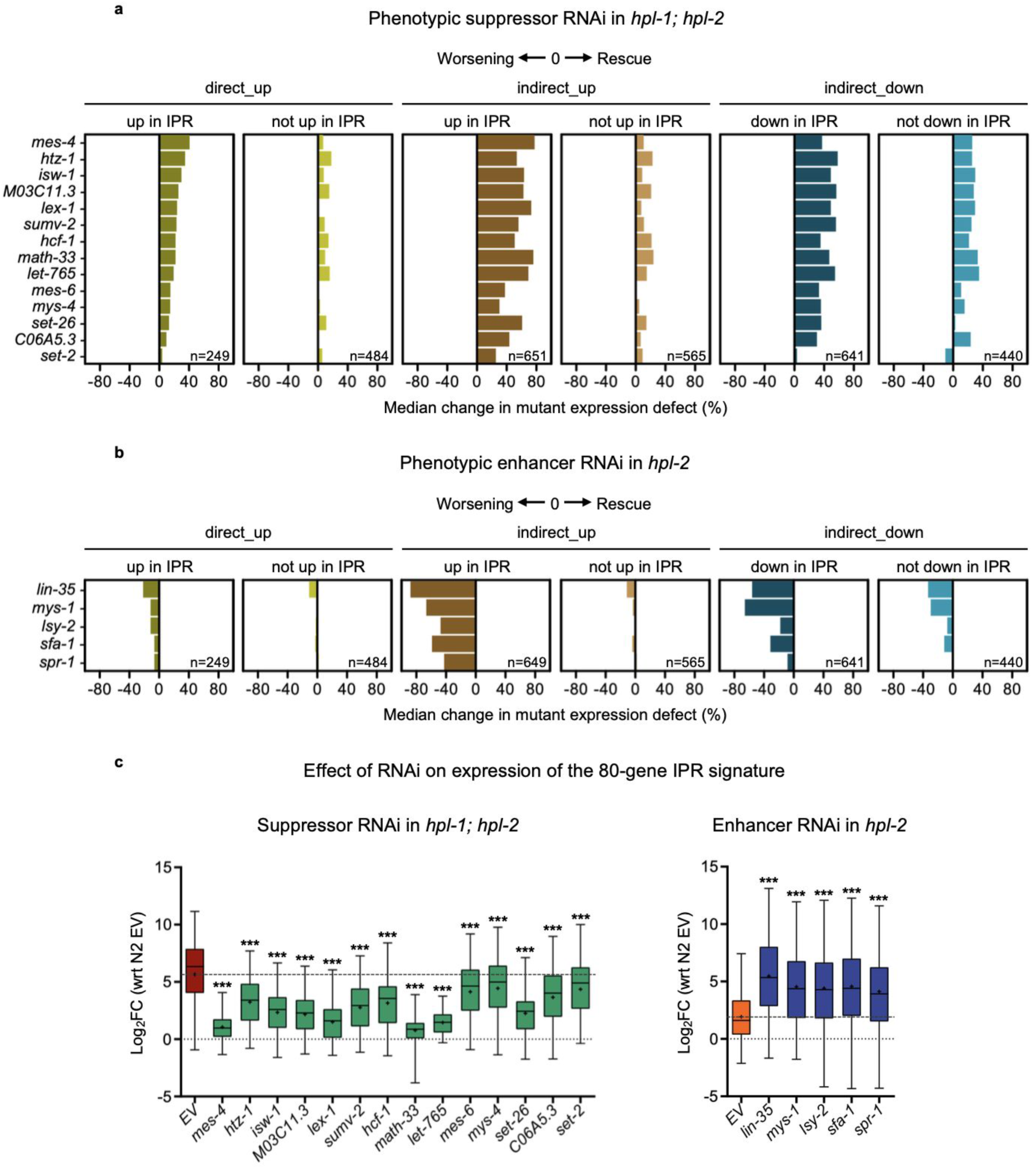
Suppressor and enhancer RNAi oppositely affect the IPR-associated transcriptional state. **a**,**b**, Median percent change in the expression of *hpl-1; hpl-2* direct_up, indirect_up, and indirect_down targets upon RNAi of (**a**) suppressors or (**b**) enhancers. Targets are further subclassified as those shared or not shared with IPR signatures. **c**, Box plots showing the effect of suppressor RNAi (left) and enhancer RNAi (right) on the expression of the 80-gene IPR signature. Two-tailed Wilcoxon signed-rank test (paired, non-parametric) was used to compare log_2_ fold changes between empty vector control (EV) and each RNAi treatment (*** Benjamini-Hochberg FDR-adjusted p-value: *p* <0.0001).

We next depleted five heterochromatin enhancers and analysed gene expression in *hpl-2* single mutants, a more suitable background to observe amplification of gene expression changes. The enhancers consisted of a member of the DRM complex (*lin-35*), a zinc finger transcription factor (*lsy-2*), and factors involved in mRNA splicing (*sfa-1*) and chromatin modifications (*spr-1, mys-1*). We observed that enhancer RNAi led to a more pronounced deregulation of indirect targets than direct targets (Fig. 3b; Extended Data Fig. 6b and Supplementary Table 10), particularly those shared with IPR pathway activation (Fig. 3b). The 80-gene IPR signature was also significantly elevated (Fig. 3c).

Thus, suppressor and enhancer depletion have opposing effects on IPR-associated transcription that mirror their effects on heterochromatin mutant growth, providing further evidence that IPR activation contributes to the growth defects. We suggest that the modest changes in direct target expression caused by suppressor and enhancer depletion alter signals that trigger the IPR, leading to the larger changes observed in the downstream IPR-associated transcriptional state.

### Derepressed transposon RNAs contribute to the heterochromatin mutant growth defects

Repression of repetitive elements is a conserved function of heterochromatin in animals^39,40^, and we previously showed that repeats are derepressed in *C. elegans* heterochromatin mutant adults^10^. We found that repetitive elements were similarly upregulated in mutant L1 larvae, with *hpl-1; hpl-2* double mutants showing the largest number (n = 339; Extended Data Fig. 7a and Supplementary Table 11), of which 75% were direct targets (Extended Data Fig. 8a–d and Supplementary Table 12). Notably, only 8% of repeats appeared to be transcribed intrinsically; the remainder lay within broader domains of elevated transcription (Extended Data Fig. 8a–d and Supplementary Table 12).

Since the IPR can be activated by viral infection, we asked whether derepression of repetitive elements might contribute to IPR activation and slow growth in heterochromatin mutants. We previously found that repetitive elements upregulated in heterochromatin mutant adults are enriched for transposase-encoding elements^10^. We observed a similar enrichment in *hpl-1; hpl-2* L1 larvae: 10.5% of transposase-encoding repeats were upregulated, compared with only 0.54% of all annotated repeat elements (Supplementary Table 12). RNAi targeting the MIRAGE1 DNA transposon was previously shown to partially restore fertility to heterochromatin mutant adults, but the effect on growth was not tested^10^.

To test whether expression of derepressed transposase ORFs contributes to the slow-growth phenotype, we performed RNAi against 16 upregulated transposase-encoding elements for which clones were available (14 direct and 2 indirect; Supplementary Table 12). We found that knockdown of MIRAGE1, CER1, Turmoil1 and NeSL-1 (all of which are in the direct class) significantly improved growth (Extended Data Fig. 7b). Notably, increased transcription of these four ORFs appears to arise from the elements themselves, whereas the remaining 12 ORFs, whose knockdown did not improve growth, lie within broader regions of elevated transcription initiated upstream of the element. These results indicate that expression of specific transposon-derived sequences contributes to the growth defects caused by heterochromatin loss and are consistent with repetitive element activation being a potential source of signals that induce the IPR.

### Mild transcriptional dampening rescues HP1 loss phenotypes in *C. elegans* and human cells

As many suppressors encode proteins associated with active chromatin, we considered that one mode of suppression might involve dampening of global transcription. Consistent with this idea, although RNAi of most RNA polymerase II (RNAPII) transcriptional components causes lethality in wild type, we observed that RNAi of three that caused mild or no defects in wild type resulted in heterochromatin phenotype suppression (Supplementary Table 2). This motivated us to directly test whether mild reduction in RNAPII activity could rescue heterochromatin phenotypes.

We selected 27 RNAPII-associated factors whose RNAi causes lethality or growth arrest in wild type and performed weak RNAi by diluting RNAi bacteria 1:9 with empty vector bacteria. Remarkably, 22 of 27 (81%) treatments significantly improved *hpl-1; hpl-2* growth (Fig. 4a and Extended Data Fig. 9a). As controls, we performed weak RNAi of essential genes in other pathways (cell cycle, mRNA processing, DNA replication) and observed reproducibly improved growth in only 2 of 18 (11%) cases (Extended Data Fig. 9b).

**Figure 4.**
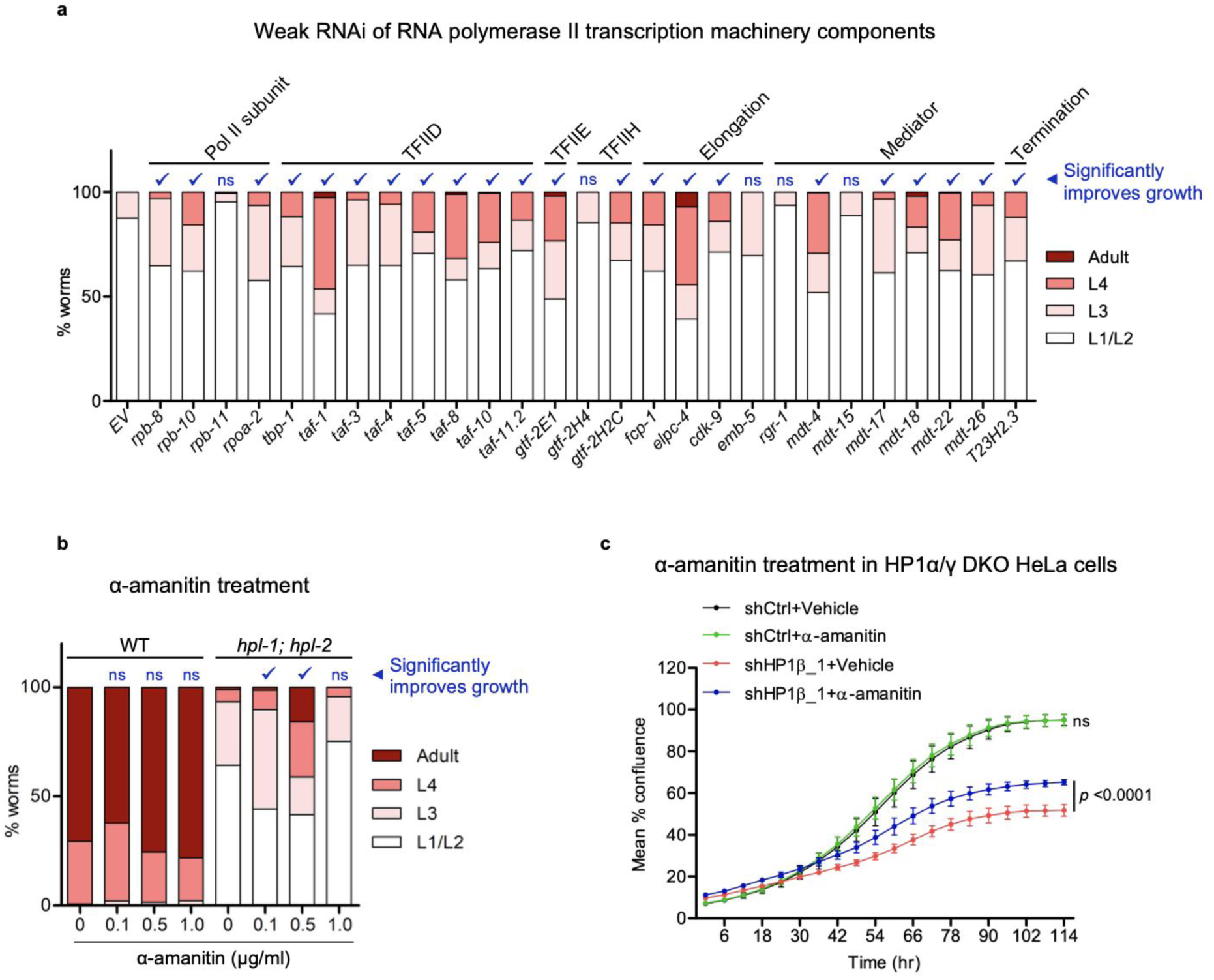
Mild inhibition of RNA polymerase II activity suppresses the growth defect of heterochromatin mutants. **a**, Developmental stages of *hpl-1; hpl-2* mutants upon weak RNAi knockdown of various components of RNA polymerase II machinery or empty vector control (EV) grown for the same length of time at 23°C. Representative data from one of two independent replicates is shown (at least 95 worms per treatment in each replicate). Chi-square tests were performed between target RNAi and control (EV) RNAi, and “✓” denotes RNAi knockdowns resulting in significantly faster growth (Bonferroni-adjusted p-value: *p* <0.05, ns: not significant). **b**, Developmental stages of wild-type (WT) or *hpl-1; hpl-2* mutants upon treatment with α-amanitin after growth for the same length of time at 23°C. Representative data from one of three independent replicates is shown (at least 190 worms per treatment in each replicate). Chi-square tests were performed between drug treatments and respective control (0 μg/ml), and “✓” denotes treatments resulting in significantly faster growth (Bonferroni-adjusted p-value: *p* <0.01, ns: not significant). **c**, Mean percent confluence of HP1α/γ double knockout (DKO) HeLa cells with (shHP1β_1) or without (shCtrl) HP1β depletion, treated with 6.25 ng/ml α-amanitin or vehicle (experimental design in Extended Data Fig. 10a). Each time point shows mean ± standard deviation from duplicate wells. Two-way mixed ANOVA was performed to assess growth trajectory divergence between vehicle and drug treatments: shCtrl+vehicle vs shCtrl+α-amanitin (F = 0.1235, *p* = 1.0) and shHP1β_1+vehicle vs shHP1β_1+α-amanitin (F = 14.81, *p* <0.0001). Bonferroni post-hoc tests were performed for pairwise differences at each time point: shHP1β_1+vehicle vs shHP1β_1+α-amanitin (*p* <0.05 from 48 hrs onwards). ns: not significant.

To directly test the effects of mildly inhibiting RNA polymerase II activity on heterochromatin growth phenotype, we treated *hpl-1; hpl-2* mutants with low doses of the inhibitor α-amanitin. We observed significantly improved *hpl-1; hpl-2* growth at concentrations that did not impair wild-type growth (Fig. 4b and Extended Data Fig. 10d), demonstrating a beneficial “dose window” of transcriptional dampening that ameliorates phenotype.

Finally, we asked if a similar relationship between heterochromatin loss and transcriptional activity may occur in humans. Heterochromatin dysfunction is implicated in human disease^41^. The human genome has three HP1 orthologs: HP1α, HP1β and HP1γ. In HeLa cells lacking HP1α and HP1γ, depletion of HP1β using two independent shRNAs slows proliferation (Fig. 4c and Extended Data Fig. 10b,c - compare black and red traces). Strikingly, low-dose α-amanitin at a concentration that does not affect HP1α(-); HP1γ(-) cells significantly improved proliferation of cells deficient in all three HP1 orthologs (Fig. 4c and Extended Data Fig. 10a-c). This suggests that transcriptional dampening can mitigate consequences of heterochromatin loss across species.

## Discussion

Heterochromatin dysfunction is associated with diverse human diseases^41^, yet the mechanisms that underlie the resulting pathological phenotypes are not well understood. Our study identifies indirect activation of the Intracellular Pathogen Response (IPR), an innate immune/stress pathway, as a major contributor to the impaired growth and development caused by heterochromatin disruption in *C. elegans*. Importantly, our finding that mild inhibition of RNA polymerase activity improves the growth of both *C. elegans* HP1 mutants and human cells lacking HP1 function suggests transcriptional dampening as a potential therapeutic avenue for diseases involving heterochromatin dysfunction.

The IPR is a multifunctional innate immune/stress pathway induced by diverse triggers, including intracellular pathogens such as viruses and fungi, proteotoxic stress, perturbation of purine metabolism and prolonged heat stress^32–35,42^. Although their components differ, the IPR and the mammalian interferon pathway share several features: they are induced by similar triggers, promote pathogen defence and can impair development^36^. Activation of the IPR causes slow development and reduced fertility, resembling phenotypes caused by loss of heterochromatin function.

The IPR has been defined by 80 genes that are strongly and commonly induced by different triggers^32–34,38^. Here, we identified the wider gene expression changes associated with IPR activation by profiling mutants of the negative regulator *pals-17*. These changes showed a high correspondence with the indirect gene expression changes induced by heterochromatin dysfunction, that is, the upregulation of genes not marked by heterochromatin and the downregulation of diverse metabolic pathways. Strikingly, knockdown of suppressor genes that improve heterochromatin mutant growth substantially corrected IPR-associated gene dysregulation, while having only mild effects on direct heterochromatin targets. Knockdown of enhancers had the opposite effect, further slowing growth and increasing IPR activation. These findings identify IPR activation as a major contributor to the physiological growth defects of heterochromatin mutants.

About 85% of the genes in our heterochromatin genetic interaction network have human orthologues, highlighting the conservation of many components of these pathways. Genes whose knockdown enhanced phenotype were enriched for those involved in ribosome biogenesis and ubiquitin-mediated proteolysis, suggesting that protein synthesis and proteostasis pathways buffer the effects of heterochromatin dysfunction. Notably, perturbations of proteostasis can induce the IPR^32,33^, suggesting that disruption of these pathways by RNAi may increase IPR activation by exacerbating proteotoxic stress.

Another class of enhancers comprised diverse chromatin regulators associated with transcriptional repression, including members of the DRM, NuRD and MEC complexes, SynMuv genes and genes in the SUMOylation pathway (Supplementary Table 2), loss of which causes slow growth^16,24,43–47^. In recent studies, upregulation of IPR reporter *pals-5*p::GFP was observed upon depletion of regulators in all of these classes, including *hpl-2, lin-61* and *met-2*^48,49^. In addition, it was shown that the 80-gene IPR signature is induced upon depletion of chromatin repressors *lin-15B, lin-35/*Rb, and *let-418/*Mi-2^48,49^. The studies proposed that the chromatin regulators play a role in directly repressing IPR genes. Our results show that the IPR is indirectly activated in heterochromatin mutants. We suggest that IPR induction upon depletion of the other chromatin repressors may also be a secondary stress response. In addition, knockdown of some of our heterochromatin suppressors was previously shown to suppress the slow growth or developmental defects of some of the chromatin repressor mutants^16,19,20,24^. We propose that this suppression may result from reduced IPR activation.

Parallels can also be seen in *Drosophila.* Disruption of the chromatin repressor L(3)MBT, an orthologue of the *C. elegans* protein LIN-61 studied here, leads to inappropriate activation of germline programmes in somatic tissues^50–53^. Through reanalysis of recent RNA-seq data from L(3)mbt-depleted ovarian somatic cells ^54^, we found that, in addition to the reported upregulation of germline genes, these cells show further similarities to *C. elegans* heterochromatin mutants: innate immune pathways are upregulated, whereas metabolic and translation-related pathways are downregulated (Supplementary Table 13). This suggests that activation of innate immune pathways and associated physiological changes may be a common consequence of defective chromatin repression.

What might trigger IPR activation in heterochromatin mutants? Repetitive element derepression is likely to make a contribution. Repetitive elements are upregulated in heterochromatin mutants, and most are direct targets of heterochromatin repression. We found that RNAi against four upregulated transposases (MIRAGE1, CER1, Turmoil1 and NeSL-1) partially improved heterochromatin mutant growth, supporting contribution from derepression of these elements. Proteotoxic stress is another plausible trigger, because translation of a large number of ectopically expressed transcripts, including germline genes expressed in the soma, could disrupt proteostasis.

In summary, we propose a model in which direct transcriptional changes following heterochromatin loss generate proteotoxic stress and transposable element-derived signals that in turn activate the IPR and potentially other stress-responsive transcriptional programs. The resulting transcriptional reprogramming would then impair metabolism and growth. Importantly, knockdown of chromatin regulators associated with active transcription, or direct inhibition of RNA polymerase II function, ameliorated heterochromatin mutant growth defects. We propose that these interventions act by modestly reducing the primary transcriptional consequences of heterochromatin loss, thereby lowering signals that induce the IPR and attenuating the downstream transcriptional response. (Fig. 5).

**Figure 5.**
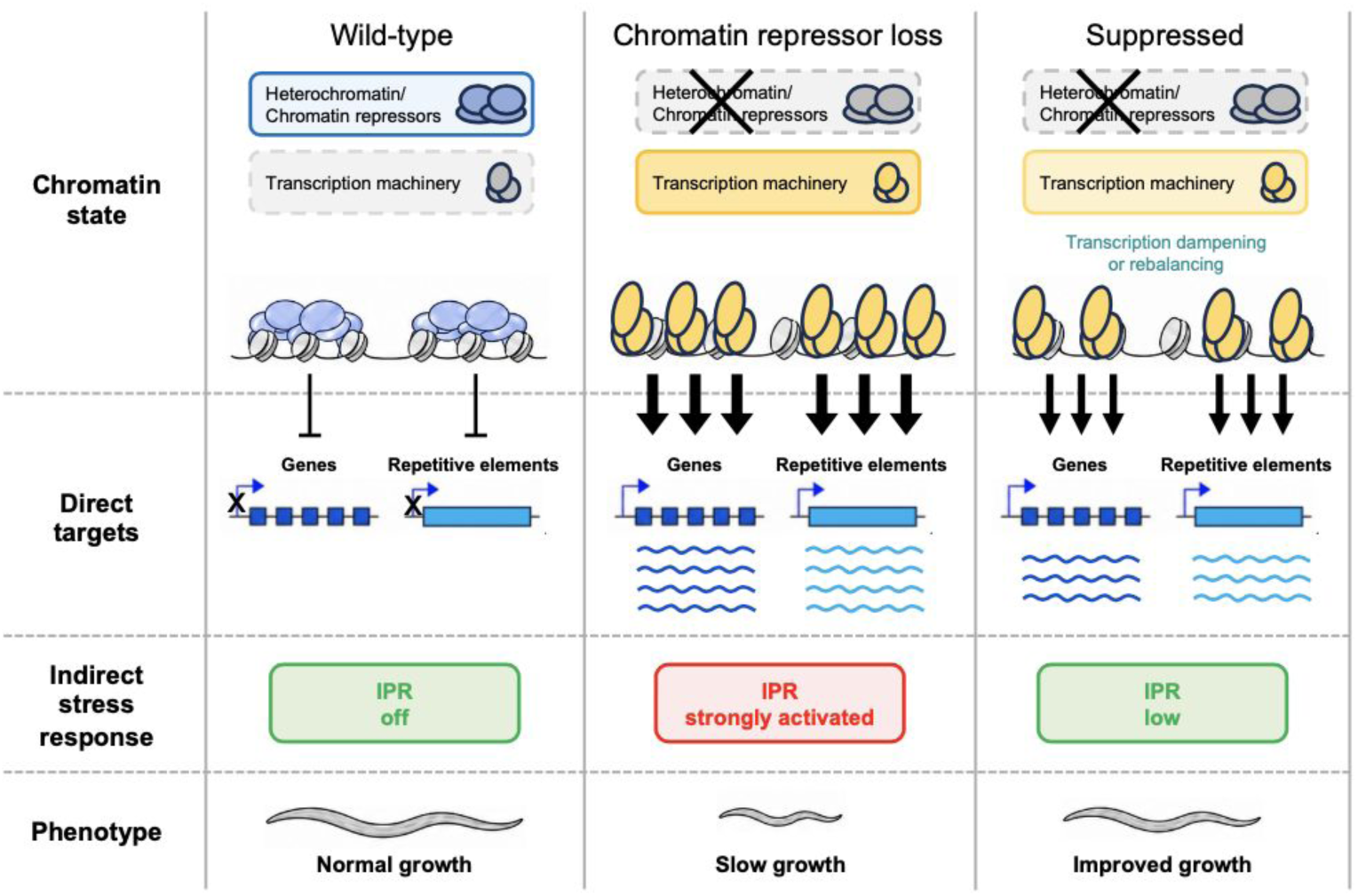
Model linking heterochromatin loss to IPR activation and growth defects. Loss of heterochromatin/chromatin repressors induces transcriptional changes, resulting in direct derepression of genes and repeats that generate proteotoxic stress and transposable element-derived signals. These in turn activate the IPR and potentially other stress-responsive transcriptional programs, leading to downstream transcriptional changes that impair metabolism and growth. Knockdown of chromatin regulators associated with active transcription, or direct inhibition of RNA polymerase II function, reduces the stress response and ameliorates growth defects in heterochromatin mutants. Reducing transcriptional output may mitigate the consequences of heterochromatin dysfunction by mildly reducing signals that induce the IPR (Figure created using Biorender.com).

Heterochromatin abnormalities are found in diverse human disorders, including cancer, neurodevelopmental disorders, premature aging syndromes and diseases caused by mutations in chromatin repressors^55–58^. Our results raise the possibility that, in at least some of these conditions, secondary immune and stress responses themselves contribute substantially to pathology. Mild reduction of RNA polymerase II activity improved the growth of *C. elegans* HP1 mutants and the proliferation of human HP1-deficient cells, indicating that heterochromatin dysfunction phenotypes can be ameliorated without restoring heterochromatin itself. We suggest that modest transcriptional dampening or targeted inhibition of conserved suppressor pathways could be explored as strategies for mitigating diseases associated with loss of chromatin repression.

## Methods

### Worm culture and strains

Strains used in this study are described in Supplementary Table 14. Worms were maintained at 15°C and cultured using standard methods^59^.

### Genetic interaction screen

We screened a sub-library of 2292 RNAi clones targeting genes encoding nuclear proteins (Supplementary Table 1; clones are from^60–62^). RNAi bacteria were grown in LB containing carbenicillin (25 µg/ml) and tetracycline (10 µg/ml) at 37°C with shaking (200 rpm) for 16–20 h, seeded onto NGM agar 6-well plates supplemented with carbenicillin (25 µg/ml) and IPTG (1 mM), and induced at room temperature for at least three days before use. An empty vector (EV; HT115 carrying L4440 without insert) was included as a non-targeting control in all experiments.

RNAi feeding experiments started with L3 worms prepared as follows: mixed-stage embryos were obtained by hypochlorite treatment of gravid adults and hatched in M9 buffer for 18–20 h at 20°C to generate synchronised starved L1 larvae (sL1s). sL1s were grown on NGM plates seeded with OP50 bacteria (∼3,000 worms per 90 mm plate) at 15°C to the mid-L3 stage (52 h for wild type; 54 h for *hpl-1*, *hpl-2*, and *lin-61*; 56 h for *met-2*; 60 h for *lin-13*; 70 h for *hpl-1; hpl-2*). Worms were washed off plates in M9 + 0.01% Triton X-100, washed, resuspended at ∼1 worm/µl in deionised water and then three worms were dispensed per well into RNAi plates using a COPAS BioSorter (Union Biometrica; settings: sort = 3, delay = 11.5, width = 9; flow rate 10–20 worms/s). Mid-L3 animals were gated based on time of flight (axial length) and extinction (optical density).

### Scoring RNAi phenotypes and genetic interactions

RNAi clones were screened in duplicate in the primary screen and in triplicate in the secondary and tertiary screens. After worm aliquoting, plates were transferred to the appropriate temperatures (*hpl-1, lin-61, met-2* enhancers at 25°C; *hpl-2* enhancers at 20°C and 25°C; *lin-13* enhancers and suppressors at 23°C; *hpl-2* and *lin-13* suppressors at 25°C). Phenotypes of P0 and F1 generations were scored on day 4 (25°C) or day 5 (20°C, 23°C) using a dissecting microscope.

Enhancers were identified by comparing RNAi phenotypes in each mutant to wild type (N2), relative to the respective empty vector (EV) control. P0 sterility was scored in three brood size categories (0–5, 6–20, 21–50), and F1 growth was assessed by developmental stage relative to wild type. Clones were advanced to the secondary screen if the primary screen showed ≥2 sterility categories or ≥2 growth stages worse than wild type. For suppressors, only growth was assessed, and clones were advanced if at least one well showed ≥1 developmental stage faster growth. All clones passing the primary screen in any mutant were tested across all mutants in the secondary screen, and the same criteria were applied for progression to the tertiary screen.

Screening results suggested mild RNAi hypersensitivity in *met-2* and *lin-61*, as the RNAi-induced growth and fertility defects were often stronger in these mutants and they appeared to have more enhancers than the other strains. To reduce false positives, only strong enhancers were retained for these mutants. In addition, *lin-13* showed variable brood size at 23°C, potentially inflating sterility scores; therefore, only strong sterility enhancers were retained for this mutant. Previous reports of RNAi hypersensitivity in these strains are inconsistent and based on limited datasets^13,15,25,63^.

Final enhancers and suppressors were defined based on reproducibility across screens. Sterility enhancers required a ≥1 category difference from N2 for *hpl-1* and *hpl-2*, and ≥2 categories for *lin-61*, *met-2*, and *lin-13*, in at least two replicates across two screening rounds. Growth enhancers required ≥1 stage delay for *hpl-1*, *hpl-2*, and *lin-13*, and ≥2 stages for *lin-61* and *met-2*, again in ≥2 replicates across two rounds (≥4/8 replicates, 50%). For these comparisons, the N2 reference was the worst phenotype observed in >2 replicates across all screens at the relevant temperature for each RNAi clone. Growth suppressors required ≥1 stage faster growth relative to EV for *hpl-2*, *lin-13*, and *hpl-1; hpl-2* under the same reproducibility criteria (Supplementary Table 2).

### Determining growth stages by DAPI staining

Synchronised starved L1s (sL1s) were obtained by hatching embryos at 15°C for 24 hours, then growing worms to L4 stage on NGM plates seeded with OP50 bacteria at 15°C and transferring L4s to the desired growth temperature - 15°C/ 20°C/ 25°C, until gravid. Embryos were then obtained by bleaching, and sL1s were hatched at the corresponding growth temperature (15°C for 24 hours, 20°C and 25°C for 20-22 hours). Unhatched embryos were removed by filtering sL1s through 20 µm Nitex. These sL1s were fed on NGM plates seeded with OP50 bacteria at corresponding growth temperatures (15°C for 96 hours, 20°C for 57 hours, 25°C for 44 hours) and then collected in M9 buffer with 0.01% Triton-X-100 (M9+Tx-100) in 1.5 ml tubes. Worms were washed once with M9+Tx-100, then pelleted at 1500 rpm for 1.5 minutes before removing supernatant and leaving ∼100 µl above the packed worm pellet. The pellet was then re-suspended in 800 µl 100% methanol and DAPI solution to final concentration 1 µg/ml and incubated for at least 30 minutes at -20°C. Following this, tubes were spun briefly at room temperature and supernatant removed, leaving ∼100 µl above the packed worm pellet. 1 ml of M9+Tx-100 was added and the tube left at room temperature for at least one hour to allow worms to rehydrate. Tubes were then spun at 1500 rpm for 1.5 minutes and supernatant removed, leaving ∼100 µl above the packed worm pellet. The pellet was washed once with 1 ml M9+Tx-100, before discarding the supernatant, leaving ∼10µl above the packed worm pellet. Worms were re-suspended in the remaining volume, mounted on a slide and growth stages counted for each sample. For statistical analysis, Chi-square test of independence (with Bonferroni correction for multiple comparisons) was used to compare growth rates between mutant and wild-type (N2) worms.

### Quantification of nucleolar size

Four synchronised L4 worms carrying the DAO-5::GFP transgene (wild type or mutant) grown from starved L1s at 15°C were transferred to fresh NGM plates seeded with OP50 bacteria and shifted to the desired growth temperature (25°C for *hpl-1, hpl-2, lin-61* and *met-2*; 23°C for *hpl-1; hpl-2* and *lin13*; 20°C for *pals-17*). Once the F1 worms reached L4 stage, 30-40 worms were picked onto 2% agarose pads in 5 mM Tetramisole for live imaging. Stage L4.4-L4.6 worms ^64^ were chosen using DIC and images in GFP channel were acquired with Volocity 6.3 on Zeiss Axioplan 2 microscope (63X Plan-Apochromat 1.4 NA oil immersion objective; Zeiss Immersol 518 F, #10539438) using Hamamatsu ORCA-ER digital camera (C4742-80). The number and area of nucleoli in the posterior hyp10 nucleus of each worm was measured using FIJI^65^. For statistical analysis, two-tailed Mann-Whitney U tests and Chi-square tests of independence were used to compare nucleolar areas and nucleolar numbers respectively, between mutant and wild-type (N2) worms. Only nuclei with a single nucleolus were considered for nucleolar area comparisons.

### Quantification of phospho-eIF2α by western blot

Synchronised starved L1s (sL1s) were obtained by hatching embryos at 15°C for 24 hours, then growing worms to L4 stage on NGM plates seeded with OP50 bacteria at 15°C and transferring L4s to 25°C until gravid. Embryos were then obtained by bleaching, and sL1s were hatched at 25°C for 20-22 hours. Unhatched embryos were removed by filtering sL1s through 20 µm Nitex. Filtered worms were then spun at 1500 rpm for 1.5 mins and supernatant removed, leaving ∼10µl above the packed worm pellet. Tubes were flash-frozen in liquid nitrogen and stored at -80°C. Protein samples were prepared by thawing worm pellets on ice in the presence of RIPA buffer (50 mM Tris-Cl pH 7.4, 150 mM NaCl, 1% Triton X-100, 0.5% Sodium deoxycholate, 0.1% SDS, 1 mM EDTA) with 1 mM PMSF and Roche protease inhibitor cocktail before sonicating at 4°C for 10 minutes (30s on/30s off; 5 minutes of active sonication) on high power (Diagenode Bioruptor Standard #UCD-200). Samples were then centrifuged (15000 rpm, 15 minutes, 4°C) and the supernatant collected. Samples were diluted with 4X Laemmli buffer and boiled at 95°C for 10 minutes before running on 4-12% NuPAGE gels in MES SDS buffer. Proteins were transferred to 0.2 µm PVDF membrane using Invitrogen iBlot dry blotting system (20V for 7 minutes) using filter pads soaked in cold Tris-Glycine buffer. After transfer, membranes were blocked in 5% BSA in 1X TBST for 1 hour before adding primary antibodies [Rabbit anti-Phospho eIF2 alpha (Ser51, CST #3597), 1:3000; Mouse anti-Paramyosin (DSHB 5-23), 1:1000] diluted in 1X TBST with 2.5% BSA and incubating overnight at 4°C. The next day, membranes were washed three times for 10 minutes in 1X TBST with shaking followed by addition of HRP-conjugated secondary antibodies [Goat anti-Rabbit IgG (Abcam ab6721), 1:80000; Rabbit anti-Mouse IgG (Abcam ab97046), 1:60000] diluted in 1X TBST with 0.5% BSA and incubating at room temperature for 60 minutes. Membranes were then washed again, three times for 10 minutes with shaking, before visualising using chemiluminescence on Bio-Rad ChemiDoc.

Quantification of p-eIF2α levels was performed using FIJI. p-eIF2α band intensity in each sample was first normalised to its respective paramyosin loading control (p-eIF2α/paramyosin ratio), following which all mutant samples were normalised to wild-type (N2). For statistical analysis, one-tailed Mann-Whitney U tests were used to determine if mutant worms exhibited higher p-eIF2α/paramyosin ratios than the wild type (N2).

### Weak RNAi of RNA polymerase II machinery

RNAi clones used are listed in Supplementary Table 1. Weak RNAi feeding plates were prepared by mixing RNAi bacteria with control empty vector (EV) RNAi bacteria at a 10% to 90% ratio. Plates were left to dry and induce at room temperature for at least 3 days. Six synchronised mid-L3 worms grown from starved L1s at 15°C were transferred to RNAi plates and shifted to 23°C for 4-5 days. Worms were then collected for growth staging by DAPI staining as described above. RNAi was considered to significantly improve *hpl-1; hpl-2* mutant growth when in both independent replicates, the overall stage distribution differed significantly from the EV control following a chi-square test of independence (with Bonferroni correction for multiple comparisons) and the proportion of L4 and adults was greater than in the EV control. Weak RNAi treatments were also performed in wild-type N2 worms, which confirmed that they had no significant effect on growth.

### RNAi of transposase ORFs

RNAi clones used for transposase RNAi knockdowns are listed in Supplementary Table 12. Six synchronised mid-L4 stage worms grown from starved L1s at 15°C were transferred to RNAi plates and shifted to 23°C until gravid. Embryos were then obtained by bleaching, and sL1s were hatched at 23°C for 20-22 hours. Unhatched embryos were removed by filtering sL1s through 20 µm Nitex, and RNAi feeding was continued in the sL1s at 23°C for 3 days. Worms were then collected for growth staging by DAPI staining as described above. RNAi was considered to significantly improve *hpl-1; hpl-2* mutant growth when in both independent replicates, the overall stage distribution differed significantly from the EV control following a chi-square test of independence (with Bonferroni correction for multiple comparisons) and the proportion of L4 and adults was greater than in the EV control.

### Treatment with α-Amanitin

50 mm NGM plates were supplemented with α-Amanitin (Sigma #A2263) in sterile dH20 by adding 500 ul of the required concentration and allowing it to dry at room temperature for 3 hours. Six synchronised mid-L3 worms grown from starved L1s at 15°C were transferred to α-amanitin-containing plates and shifted to 23°C for 4-5 days. Worms were then collected for growth staging by DAPI staining as described above. α-amanitin treatment was considered to significantly improve *hpl-1; hpl-2* mutant growth when in both independent replicates, the overall stage distribution differed significantly from the no drug control following a chi-square test of independence (with Bonferroni correction for multiple comparisons) and the proportion of L4 and adults was greater than in the no drug control.

### Ribosomal RNA quantification

Starved L1s were grown as described in the RNA sequencing section. Frozen worm pellets were thawed in 500 ul TriPure Buffer (Roche #11667157001) and ∼250 ul 0.7 mm Zirconia beads (BioSpec Products Cat. No. 11079107 zx) were added to each sample. Samples were homogenised in Mixer Mill MM400 (2 rounds – frequency 30 (1/s) for 2 mins 30 secs, with a 1 minute cooling time between rounds). Following this, 100 ul chloroform was added and the sample shaken for 15 seconds and incubated at room temperature for 10 minutes before centrifuging for 15 minutes at 12000xg at 4°C. The aqueous phase was transferred to a new tube and 1 µl GlycoBlue (Invitrogen #AM9515) added to visualise the pellet, before combining with an equal volume of isopropanol by inverting the tube. The tube was incubated at room temperature for 10 minutes, then centrifuged for 15 minutes at 16000xg at 4°C. The pellet was washed with 1 ml 75% ethanol and all traces of ethanol removed before air-drying for 5-10 minutes. RNA pellets were resuspended in nuclease-free water to a concentration of 250 worms/ul. Ribosomal RNA concentration was measured using RNA ScreenTape (Agilent #5067-5576) and the per worm molarity of 18S ribosomal RNA was calculated for each sample using values obtained from the TapeStation Analysis Software 4.1.1. For statistical analysis, two-tailed Mann-Whitney U test was used to compare 18S rRNA levels between mutant and wild-type (N2) worms.

### RNA sequencing

RNA-seq experiments were carried out using RNA extracts prepared from samples of 1000-10,000 starved L1s (sL1s) in two independent biological replicates. Starved L1 samples for RNA extraction were prepared by first hatching synchronised sL1s at 15°C for 24 hours, then growing to L4 stage on NGM plates seeded with OP50 bacteria at 15°C and transferring L4s to the desired growth temperature - 15°C/ 20°C/ 25°C, until gravid. Embryos were then obtained by bleaching, and sL1s were hatched at the corresponding growth temperature (15°C for 24 hours, 20°C and 25°C for 20-22 hours). Unhatched embryos were removed by filtering sL1s through 20 µm Nitrex. sL1s were counted, then pelleted and buffer removed before flash-freezing in liquid nitrogen and storing at -80°C. RNA samples for the enhancer and suppressor knockdown experiments were obtained in the same manner, except that worms were grown on NGM plates seeded with OP50 bacteria at 15°C until mid-L3 stage and then transferred to HT115 RNAi-feeding plates at 25°C (∼1200 worms per 90 mm RNAi plate; RNAi plates were prepared as described above).

1 ml TriPure Buffer (Roche #11667157001) was added to frozen worm pellets before homogenising by passing the sample ten times through a 25G needle (Camlab #1150831) attached to a 2 ml syringe. Samples were then vortexed with 50-100 µl acid-washed sand (Sigma #274739) for 15 minutes at room temperature. 200 µl chloroform was then added, the sample shaken for 15 seconds and incubated at room temperature for 10 minutes before centrifuging for 15 minutes at 12000xg at 4°C. The aqueous phase was transferred to a new tube and 1 µl GlycoBlue (Invitrogen #AM9515) added to visualise the pellet, before combining with an equal volume of isopropanol by inverting the tube. The tube was incubated at room temperature for 10 minutes, then centrifuged for 15 minutes at 16000xg at 4°C. The pellet was washed with 1 ml 75% ethanol and all traces of ethanol removed before air-drying for 5-10 minutes and re-suspension in 30 µl nuclease-free water, followed by incubation for 10 minutes at 55°C. The RNA concentration and quality was measured using Qubit fluorometer RNA High Sensitivity kit and High Sensitivity RNA ScreenTape (Agilent #5067-5579) respectively, with samples showing RNA integrity number (RIN) above 9 taken forward for library preparation.

Total RNA-seq libraries were prepared from 100 to 500 ng RNA. First, genomic DNA was removed by digestion for 10 minutes at 37°C with 2U DNase I (ThermoFisher Scientific #AM2238) with 2 µl RNase inhibitor (Cambio #RG90910K) in a total volume of 50 µl. RNA was then purified using 2.2X RNACleanXP beads (Beckman Coulter #A63987), according to manufacturer’s instructions, and re-suspended in 7.5 µl nuclease-free water. Ribosomal RNAs were depleted by hybridising custom-designed DNA oligo probes to RNA samples, then degrading DNA/RNA hybrids using RNaseH as in^66^. Lastly, after RNaseH digestion of ribosomal RNA, a further DNase I digestion was carried out. rRNA-depleted RNA was then purified using 2.2X RNACleanXP beads, as before.

RNA-seq libraries were prepared from rRNA-depleted RNA using the NEBNext Ultra II Directional RNA Library Prep Kit for Illumina (New England Biolabs #E7760L), according to the manufacturer’s protocol. For purification steps, SPRIselect beads (Beckman Coulter #B23318) were used. cDNA library quality was assessed using D1000 DNA ScreenTape (Agilent, 5067-5582) and concentration measured by Qubit fluorometer dsDNA High Sensitivity kit (Invitrogen #Q32851). Sequencing was carried out using the Illumina NovaSeq platform with paired end 50 bp reads.

### RNA-Seq processing (genes and repeats)

RNAseq reads were processed using the nf-core/rnaseq v3.14.0^67^ pipeline implemented in Nextflow v24.04.2^68^. Samples split into two lanes of the same run (L001 and L002) are merged at the start. Quality control and read trimming was performed with Trim Galore v0.6.7^69^. After this, Kallisto v0.48.0^70^ pseudomapping was performed. For gene analyses, pseudomapping was run against the full “canonical geneset” (c_elegans.PRJNA13758.WS285.canonical_geneset.gtf) annotation available from Wormbase PRJNA13758 WS285 with default parameters except for ‘--rf-stranded -b 100 --genomebam’. Scaled gene counts were produced by tximport (bioconductor-tximeta: 1.12.0)^71^, rounded to the nearest integer and subsequently filtered for genes corresponding to “lincRNA”, “pseudogenes” and “protein_coding” in the canonical geneset annotation. For the repeat analyses, Kallisto pseudomapping was performed with default parameters except for ‘-b 100 --genomebam’ against unstranded Dfam2.0^72^ repeat annotation, which was further processed to remove the mitochondrial chromosome and rRNA gene annotations. Repeat counts were produced by tximport (bioconductor-tximeta: 1.12.0) and rounded to the nearest integer. Technical replicates (samples that were re-sequenced) were aggregated using DESeq2 (v1.48.1) collapseReplicates^73^. DESeq2 (v1.48.1) was used to compute differential expression (p-adj <0.001).

### ChIP-seq library preparation and data processing

Nematode growth and ChIP-seq was performed on two independent biological replicates of starved L1 larvae using the same methods as in^74^ and the same antibodies as in^10^. For H3K9me2, replicate 1 was done here and replicate 2 is sample GSM4697043 from GEO accession GSE155189^74^. ChIP-seq reads were trimmed to remove adapters, low-quality bases, and ambiguous bases (N) using Trim Galore. For paired-end samples, trimming was performed in paired mode. Trimmed reads were aligned to the *C. elegans* ce11 reference genome using BWA-MEM^75^. Alignments were coordinate-sorted and indexed with SAMtools^76,77^. Genome coverage tracks were generated using bamCoverage from the deepTools suite^78^, with reads from paired samples extended to fragment length and single end to 140bp, a bin size of 1 bp, and normalisation by counts per million (CPM); reads were excluded from ENCODE blacklist regions ^79^. Broad peaks were called for each sample using MACS2^80^ (v2.2.9.1) from alignments with MAPQ ≥10, the ‘--broad’ flag, and a broad cutoff of 0.1 using 20 million reads from GSM2333111 (GEO accession GSE87524^10^) as the input control. For H3K9me2, peak calling was also performed without imposing a minimum mapping quality threshold, thereby retaining alignments with MAPQ ≥0. Peak sets were defined as regions called in both replicates using BEDTools intersect^81^ (Supplementary Table 4).

### Target classification (direct_up, indirect_up, indirect_down)

To classify differentially expressed genes in *hpl-1; hpl-2* mutants as direct or indirect targets, BEDTools intersect was used to intersect H3K9me2 peaks (mapq10 and mapq0) with gene promoter annotations from^82^ supplemented with 500 bp upstream of the Wormbase TSS where no annotated promoter was available for the gene; promoters for the first gene in an operon were also assigned to downstream genes (Supplementary Table 4). Direct-up targets were defined as significantly upregulated genes (adjusted p-value <0.001 and log2FC > 0) in *hpl-1; hpl-2* that have a promoter marked by H3K9me2; indirect_up targets are significantly upregulated genes with no H3K9me2 peak at their promoter(s) (Supplementary Table 5). Significantly downregulated genes (adjusted p-value <0.001 and log2FC < 0) were considered indirect (indirect_down).

Repeat classification as direct or indirect targets of *hpl-1; hpl-2* was done manually. Upregulated repeats were defined as those with a log2FC > 0 and adjusted p-value <0.001, that did not overlap with introns of genes upregulated in *hpl-1; hpl-2* (Supplementary table 11). For each significantly upregulated repeat element in *hpl-1; hpl-2* mutant sL1s, stranded transcriptional signal from CPM normalised RNA-seq bigwig files was used to visually define a 500bp region encompassing the genomic region from which transcription initiated. Transcription initiation regions were also annotated for the presence of a repeat through overlap with an annotation from Dfam2.0, and/or a promoter annotation from^82^. Direct_up repeats were defined as those where the transcription initiation region overlapped an H3K9me2 peak called using mapq10 or mapq0 data, and the remainder were classified as indirect_up (Supplementary Table 12).

### Gene expression tissue specificity

Single-cell RNA-seq data for L2 larvae^83^ was used to determine tissue specificity of deregulated genes, using cell type expression for neurons, hypodermis, pharynx, body wall muscle, glia and intestine from Table S3 and somatic gonad and germ line expression from Table S4. We calculated the Coefficient of Variation (CV) of each gene (in TPMs - transcripts per million) across the eight tissues. Tissue specificity of gene expression was classified according to the following rules: first, genes with TPM<10 in every tissue were designated as “Low expression”; the remaining genes were then classified as “Ubiquitous” (CV ≤1), “Mixed” (1 < CV < 2) or “Tissue specific” (CV ≥2). For genes classified as Tissue specific, the assigned tissue was that corresponding to the tissue with maximum expression (Supplementary Table 6).

### Calculation of change in mutant gene expression defects after suppressor and enhancer RNAi treatments

To calculate the magnitude of change in *hpl-1; hpl-2* and *hpl-2* gene expression defects by suppressor and enhancer RNAi treatments, we compared gene expression changes of the mutants to changes after RNAi treatment, both relative to wild type empty vector RNAi. We used DESeq2 to compute the gene expression changes for all the comparisons (Supplementary Table 8, 10), then divided genes into three groups according to the defined *hpl1; hpl-2* targets (direct-up, indirect_up or indirect_down; Supplementary Table 5) and calculated the percentage change for all genes in each group upon different RNAi treatments as follows:

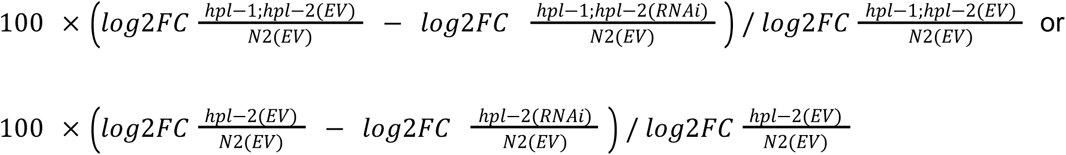

Median percentage change upon different RNAi treatments for each group was then calculated and plotted.

### Cell culture

HEK293ET and HeLa cells were a kind gift from Prof. Paul Lehner (CITIID, University of Cambridge, UK). HeLa cells were cultured in RPMI supplemented with 10% fetal bovine and penicillin/streptomycin (100 U/ml). HEK293ET cells were cultured in IMDM supplemented with 10% fetal bovine and penicillin/streptomycin (100 U/ml). Cell lines were maintained at 37°C, 5% CO_2_ and routinely tested for mycoplasma contamination (MycoBlue Mycoplasma Detector, Neo Biotech).

### Lentiviral packaging and transduction

HEK293ET cells were transfected with the lentiviral shRNA transfer plasmid, plus four packaging plasmids encoding Gag-Pol, Rev, Tat and VSV-G, using PolyJet In Vitro DNA Transfection Reagent (SignaGen Laboratories). Lentiviral supernatant was collected 48 hours post-transfection and filtered through a 0.45-µm filter before being used to transduce HeLa cells. At 48 hours post-transduction, HeLa cells were treated with 2 µg/ml puromycin for 4 days to select for shRNA expression. The shRNA hairpin sequences are as follows: shControl-GATCCACTACCGTTGTTATAGGTGTTCAAGAGACACCTATAACAACGGTAGTTTTTTTG, shHP1β_1-GATCCGCGCAAAGCTGATTCTGATTTTCAAGAGAAATCAGAATCAGCT, shHP1β_2-GATCCGTGCCCACAGGTTGTCATATTTCAAGAGAATATGACAACCTGT.

### α-Amanitin treatment and live-cell imaging

HP1α/γ double-knockout cells^84^ were transduced with shRNA vectors in duplicate (shHP1β_1 and shHP1β_2 independently), and selected with puromycin as above. After selection, cells were reseeded in 24-well plates at 20,000 cells/well. Cells were allowed to reattach overnight and then treated with 6.25 ng/ml α-amanitin in sterile water or vehicle control (water). This concentration was chosen from a dose-response survival curve (range: 3.25-100 ng/ml) because concentrations ≥12.5 ng/ml resulted in cell inviability.

Live-cell imaging was then initiated using the Incucyte S3 Live-Cell Analysis System (Sartorius). Brightfield images were taken every 6 hours for a duration of up to 114 hours of α-amanitin treatment. Incucyte analysis software was used to mask cells and compute confluency. For statistical analysis, two-way mixed ANOVA was used to compare vehicle- and α-amanitin-treated cells for both shRNAs independently. Drug treatment (vehicle vs 6.25 ng/ml α-amanitin) was modelled as a ‘between-subjects’ factor, and time was modelled as a ‘within-subjects’ factor, with individual wells serving as subjects (n=2 per treatment group). The primary term of interest was the Drug × Time interaction, which tests whether the growth trajectories of vehicle- and drug-treated cells diverge over time. Pairwise comparisons between vehicle and α-amanitin at each timepoint were performed using Bonferroni post-hoc tests.

### HeLa cell western blotting

Cells were washed once in PBS and then lysed in TBS with 1% SDS and 1:200 benzonase for 20 minutes at room temperature. Lysates were then denatured by adding Laemmli buffer and heating at 70°C for 10 minutes. Proteins were separated by SDS-PAGE and transferred onto PVDF membranes using a Trans-Blot SD Semi-Dry Transfer Cell (Bio-Rad). Membranes were blocked in 5% milk in PBS with 0.2% Tween-20 and probed with primary antibodies at 4°C overnight [Rabbit anti-HP1α (CST #2616S); Mouse anti-HP1β (Active Motif #39979); Rabbit anti-HP1γ (Abcam ab10480); Mouse anti-β-actin (Sigma A2228)].

Membranes were rinsed thrice with a wash buffer (PBS + 0.2% Tween-20) and then incubated with HRP-conjugated secondary antibodies for 40 minutes at room temperature. Membranes were then washed three more times, and chemiluminescent bands were developed on film using Pierce ECL Plus or SuperSignal West Pico Western Blotting Substrates (Thermo Fisher Scientific).

## Acknowledgements

We thank lab members for discussions and Toby Buttress and Ser van der Burght for helpful comments on the manuscript. We are also grateful to the Gurdon media team for media preparation.

## Contributions

J.A., R.P., and A.F.T. conceived the study. R.P, A.F.T, J.M.D., Y.D., Y.F., A.A., S.H., and H.V collected the data. R.P., A.F.T., A.V.P., J.M.D., F.N.C., I.T., and J.A. analysed the data. J.A., R.P, and A.F.T. wrote the original draft. All authors edited the paper and approved the final version.

---

**Extended Data Figure 1.**
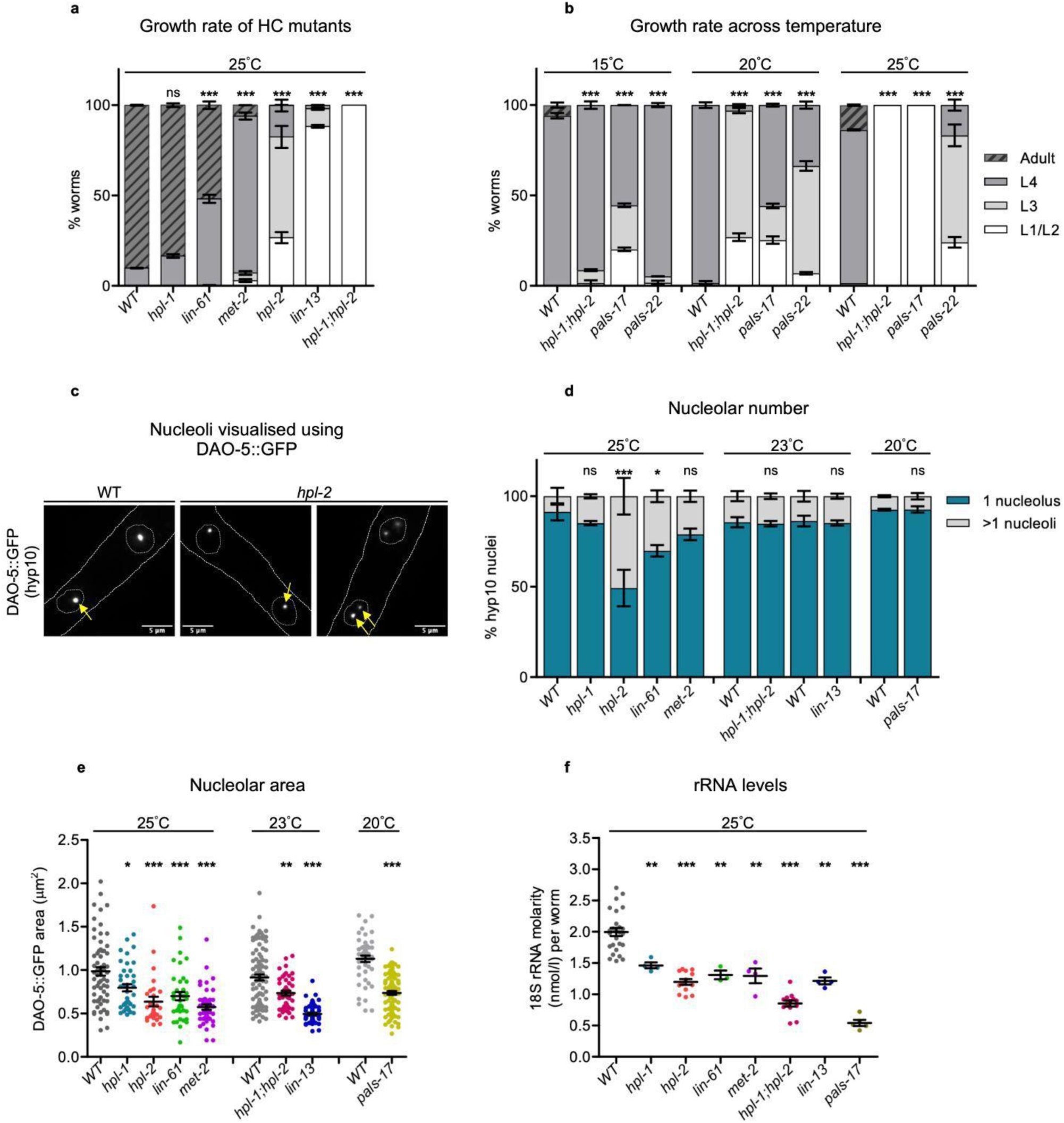
Growth rates and nucleolar phenotypes of heterochromatin and *pals* mutants. **a**,**b**, Developmental stages of wild-type (WT) and indicated mutants grown for the same length of time at (**a**) 25°C or (**b**) different temperatures. Combined data from two independent replicates is shown (at least 100 worms per genotype in each replicate). **c**, Representative fluorescence images of L4-stage wild-type (WT) and *hpl-2* worms expressing DAO-5::GFP, marking the nucleolus in hyp10 cells. Dashed lines demarcate the worm body and hyp10 nuclei. Yellow arrows point to posterior hyp10 nuclei with (i) a single nucleolus seen in most WT, and (ii) a single nucleolus or >1 nucleoli seen in *hpl-2*. Scale bar – 5 μm. **d**, Number of posterior hyp10 nuclei in L4-stage worms showing 1 nucleolus or >1 nucleoli. Combined data from two independent replicates is shown (10-20 worms per genotype in each replicate). **a**,**b**,**d**, Error bars: mean ± SEM. Chi-square tests were performed between mutants and respective WT (Bonferroni-adjusted p-value: ** *p* <0.05, *** *p* <0.0001, ns: not significant). **e**, Quantification of DAO-5::GFP area as a proxy for nucleolar size in posterior hyp10 nuclei of L4-stage worms. Combined data from two independent replicates is shown; each dot represents a single nucleolus (10-20 worms per genotype in each replicate). **f**, Levels of 18S rRNA in starved L1s. Combined data from at least three independent replicates is shown; each dot represents an independent measurement. **e**,**f**, Error bars: mean ± SEM. Two-tailed Mann-Whitney U tests were performed between mutants and respective WT (*** *p* <0.001, ** *p* <0.01, * *p* <0.05).

**Extended Data Figure 2.**
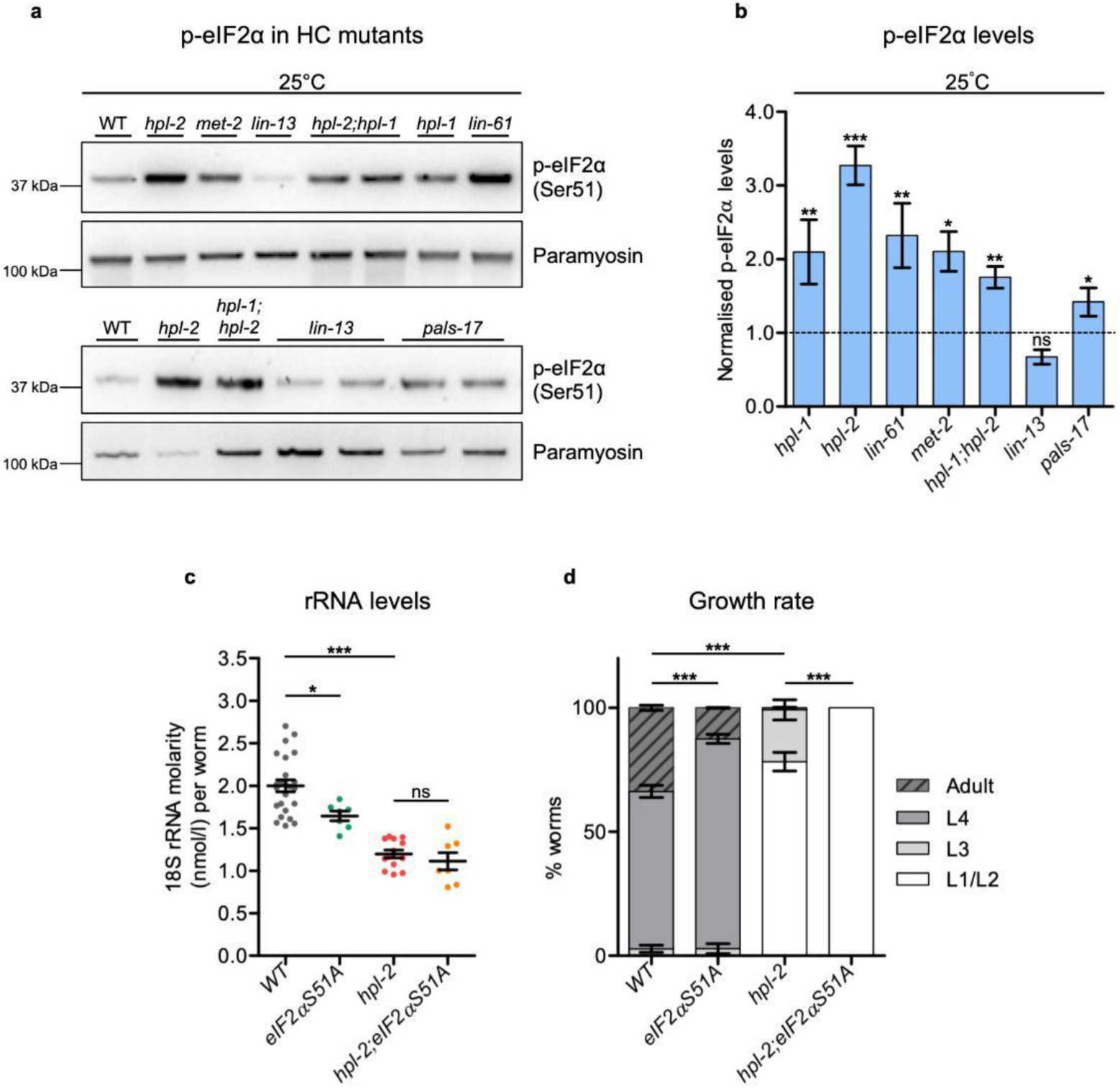
Heterochromatin mutants activate the Integrated Stress Response (ISR), which is protective. **a**, Representative western blots showing levels of phosphorylated eIF2α (p-eIF2α, Ser51) in mutant and wild-type (WT) starved L1-stage worms. Paramyosin served as a loading control. **b**, Normalised level of phosphorylated eIF2α (p-eIF2α) in mutants relative to wild type based on quantification of western blot signals in starved L1s. Combined data from at least three independent replicates; WT p-eIF2α band intensity = 1.0 [dashed line]. Error bars: mean ± SEM. One-tailed Mann-Whitney U tests were performed between mutants and wild type (*** *p* <0.001, ** *p* <0.01, * *p* <0.05, ns: not significant). **c**, Levels of 18S rRNA in starved L1-stage *hpl-2; eIF2αS51A* double mutants (25°C). Combined data from at least three independent replicates is shown; each dot represents an independent measurement. Error bars: mean ± SEM. Two-tailed Mann-Whitney U tests were performed between mutants and wild type (*** *p* <0.001, * *p* <0.05, ns: not significant). **d**, Developmental stages of wild-type (WT) and *hpl-2; eIF2αS51A* double mutants grown for the same length of time at 25°C. Combined data from two independent replicates is shown (at least 100 worms per genotype in each replicate). Error bars: mean ± SEM. Chi-square tests were performed between genotypes as indicated (*** Bonferroni-adjusted p-value: *p* <0.0001).

**Extended Data Figure 3.**
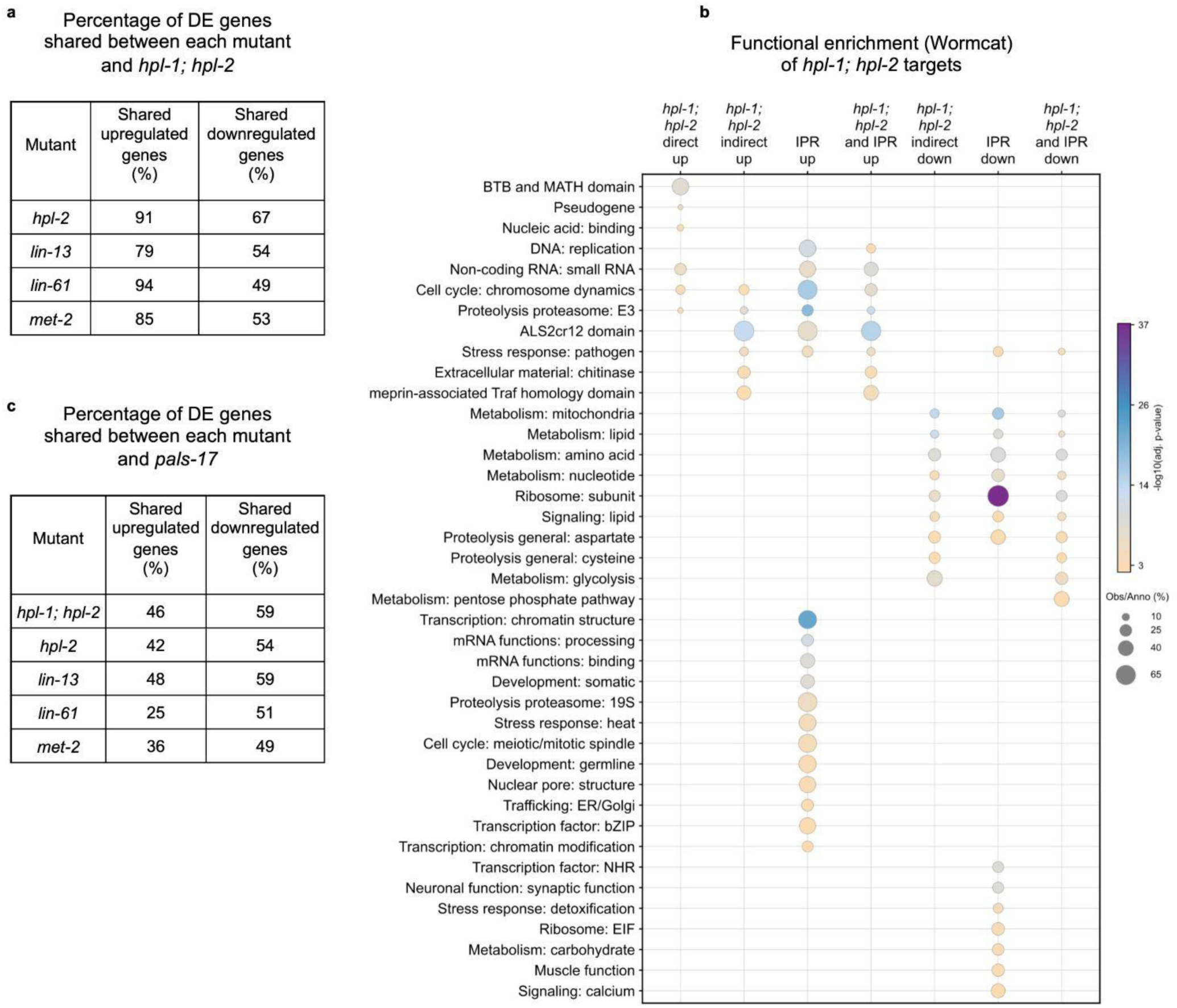
Differentially expressed gene overlap and functional enrichment in heterochromatin mutants. **a**, Percentage of differentially expressed (DE) genes shared between each heterochromatin mutant (*hpl-2, lin-13, lin-61* and *met-2*) and *hpl-1; hpl-2*. **b**, Functional category enrichment of different gene sets performed using Wormcat annotation (p-values: Fisher exact test with Bonferroni correction for multiple testing). **c**, Percentage of differentially expressed (DE) genes shared between each heterochromatin mutant (*hpl-1; hpl-2*, *hpl-2, lin-13, lin-61* and *met-2*) and *pals-17*.

**Extended Data Figure 4.**
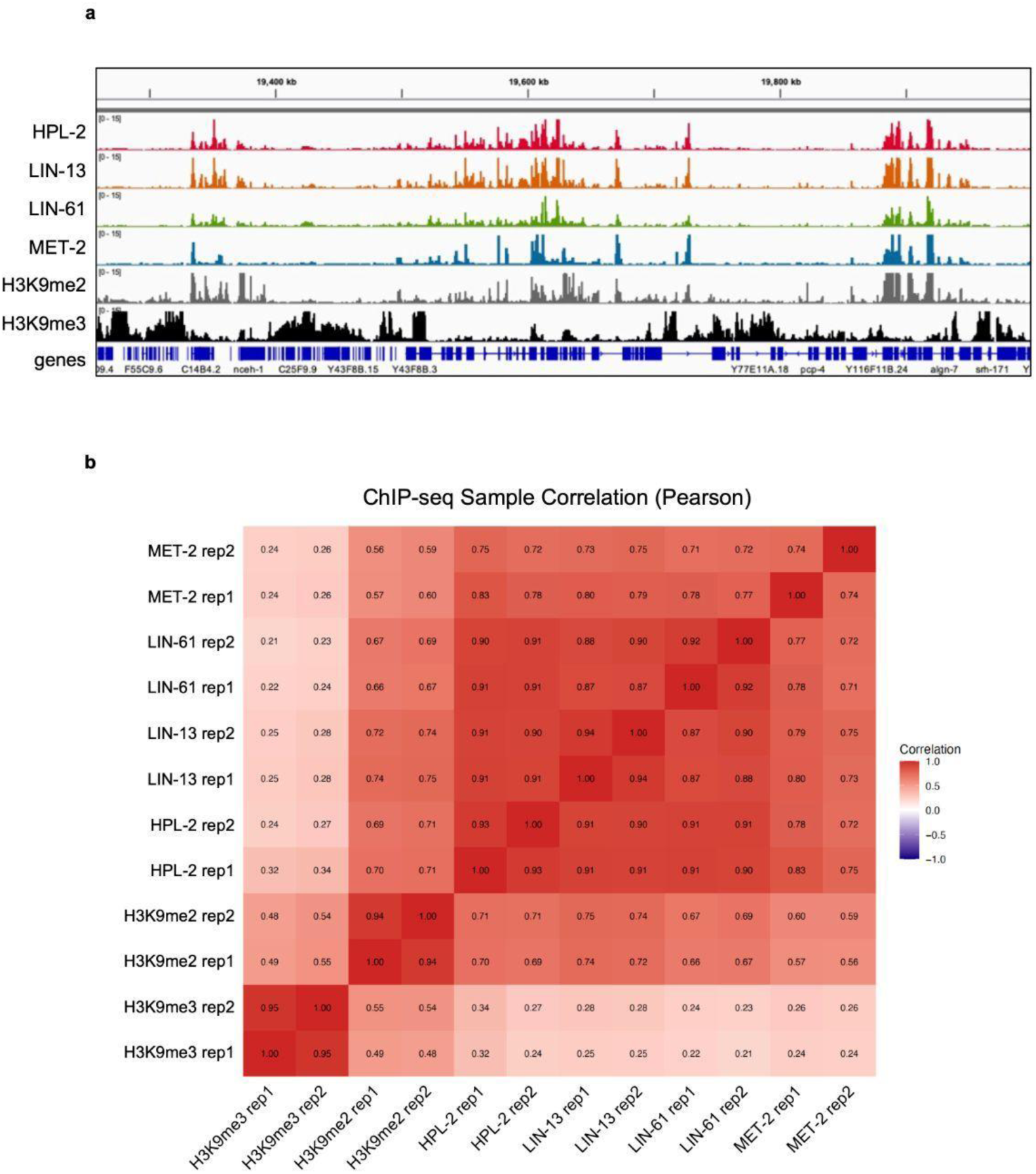
Visualisation of ChIP-seq data and correlation between datasets. **a**, IGV browser screenshot showing ChIP-seq coverage (in CPM) for the indicated heterochromatin proteins and histone modifications in starved L1 stage worms. **b**, Pearson correlation plot for all ChIP-seq samples using 1kb bins.

**Extended Data Figure 5.**
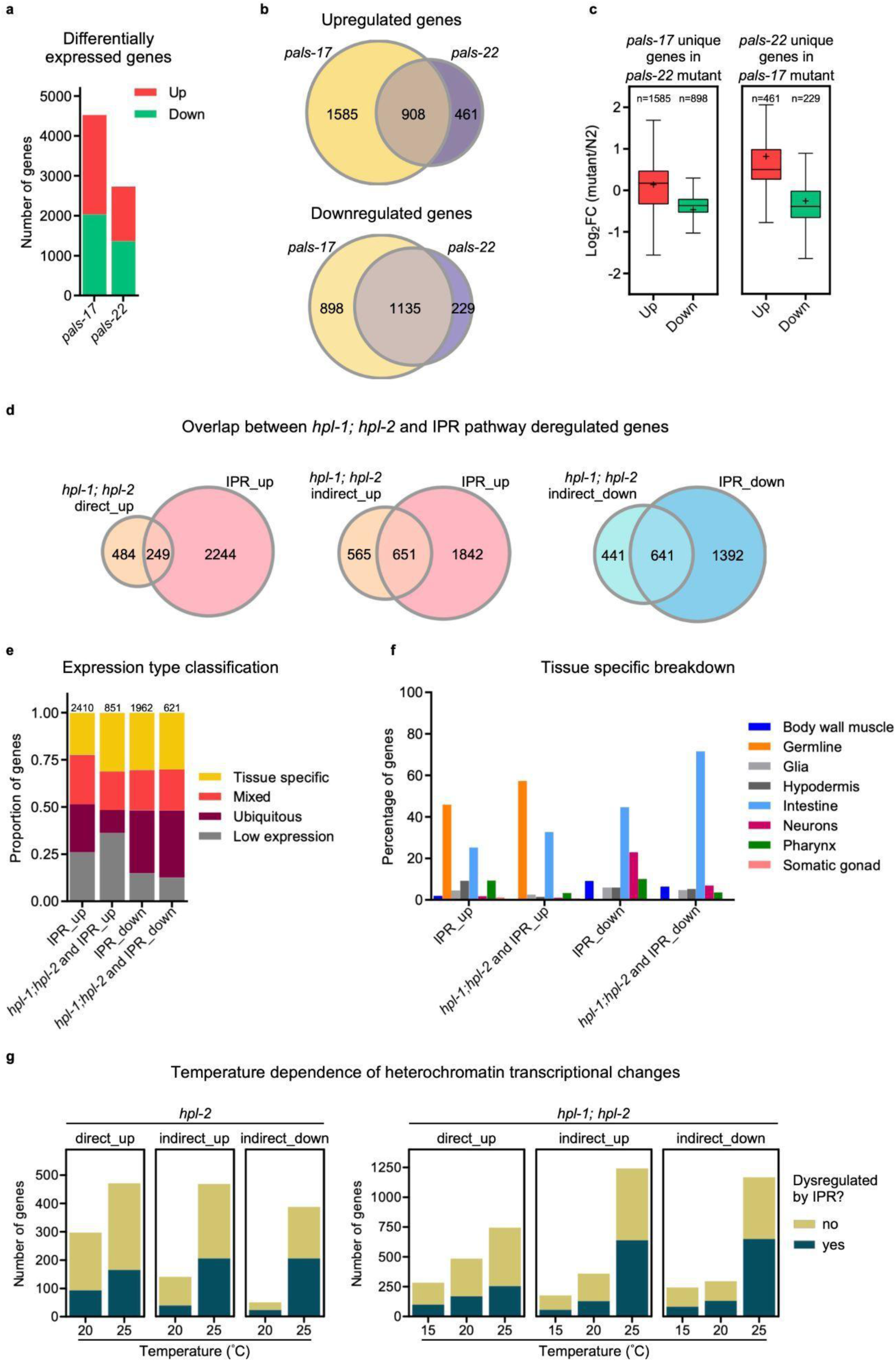
Comparison of gene expression changes in IPR mutants and the relationships between genes deregulated in heterochromatin mutants and those altered upon IPR activation. **a**, Number of up- and downregulated genes in *pals-17* and *pals-22* mutants (adjusted p-value: *p* <0.001). **b**, Venn diagram showing the extent of overlap between *pals-17* and *pals-22* mutants for up- (top) and downregulated genes (bottom). **c**, Boxplots showing the extent of deregulation of (left) *pals-17*-only differentially expressed genes in the *pals-22* mutant and (right) *pals-22*-only differentially expressed genes in the *pals-17* mutant. “only” denotes genes uniquely deregulated in one mutant and not the other. **d**, Venn diagrams showing the overlap between *hpl-1; hpl-2* direct_up genes and IPR_up genes (left), *hpl-1; hpl-2* indirect_up genes and IPR_up genes (middle), and *hpl-1; hpl-2* indirect_down genes and IPR_down genes (right). **e**, Classification of indicated gene groups into different expression types – tissue specific, mixed, ubiquitous and low expression. **f**, Tissue of maximum expression for tissue specific genes from (**e**). **g**, Number of direct_up, indirect_up and indirect_down genes at different temperatures in *hpl-2* (20°C and 25°C) and *hpl-1; hpl-2* (15°C, 20°C and 25°C) mutants, further subclassified as those dysregulated upon IPR activation at 25°C or not.

**Extended Data Figure 6.**
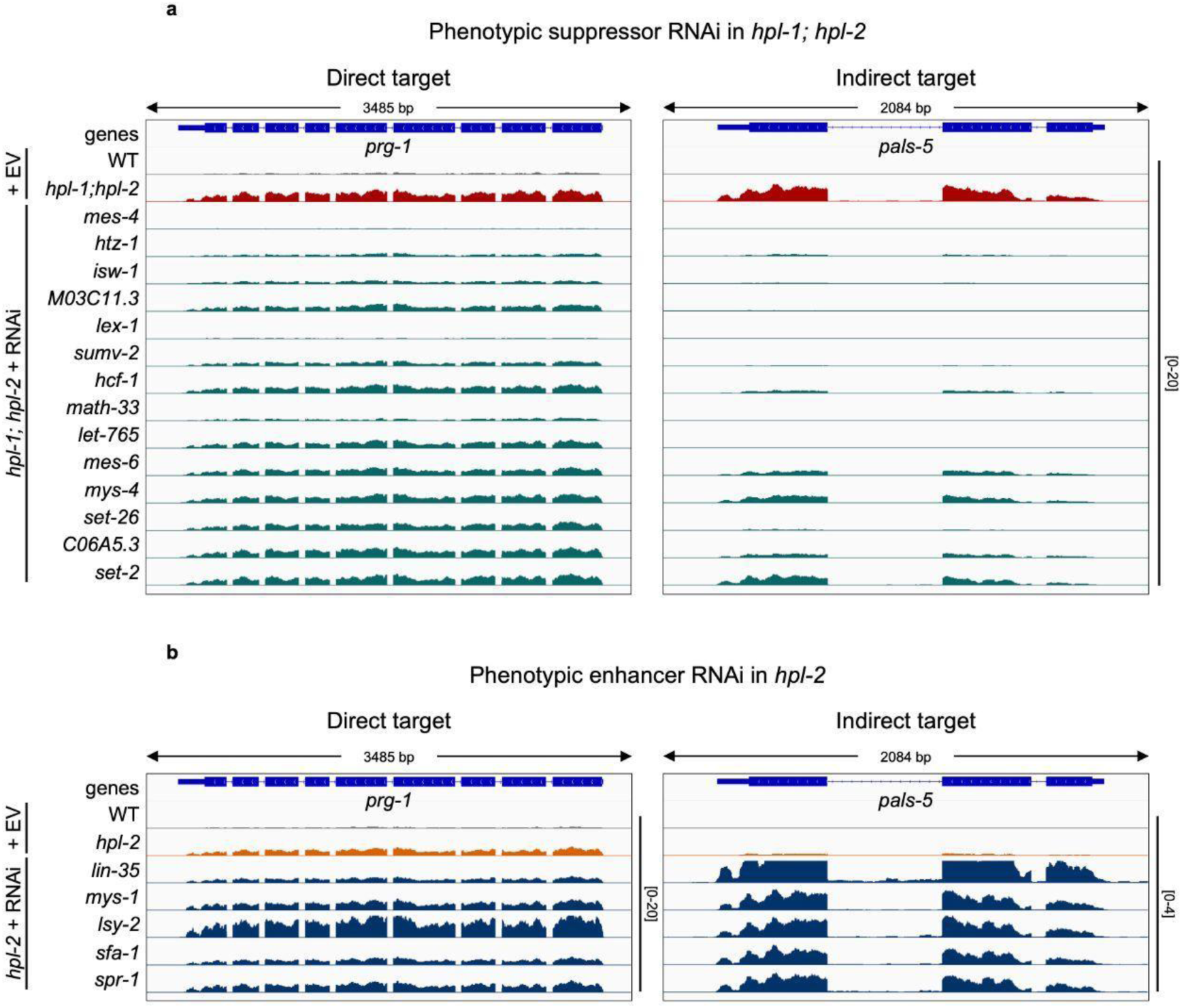
RNAi of heterochromatin suppressors reduces, while RNAi of enhancers increases, IPR-associated transcriptional changes. **a**,**b**, IGV browser screenshots of RNA-seq profiles (in CPM) for a direct target (*prg-1*) and an indirect target (*pals-5*) upon RNAi of 14 suppressors or EV (empty vector) in *hpl-1; hpl-2* double mutants (**a**) and upon RNAi of five enhancers or EV (empty vector) in *hpl-2* single mutants (**b**).

**Extended Data Figure 7.**
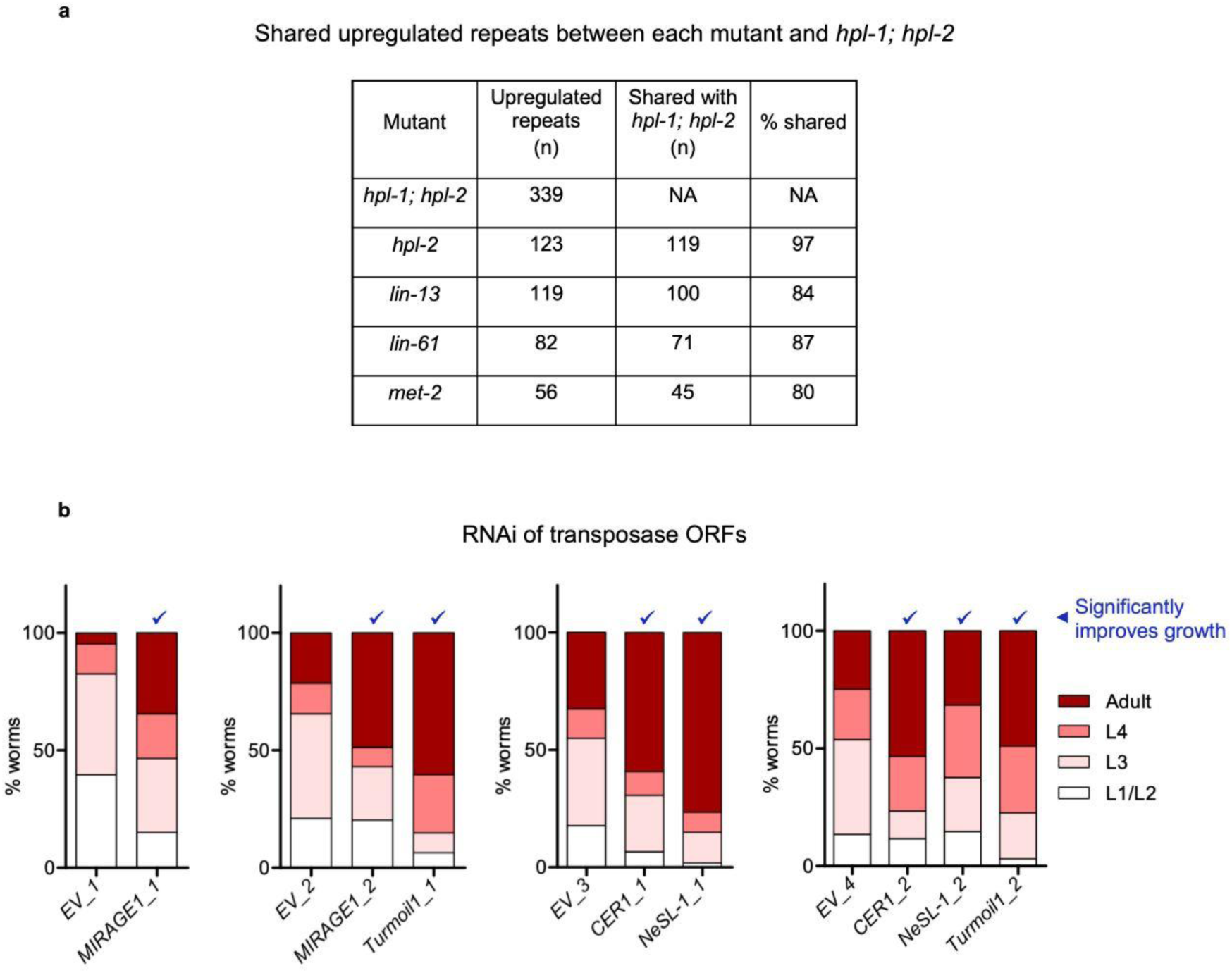
Repetitive elements are upregulated in heterochromatin mutants and transposase expression contributes to the slow growth of *hpl-1; hpl-2* mutants. **a**, Number and percentage of upregulated repeats (adjusted p-value: *p* <0.001) shared between *hpl-1; hpl-2* and the indicated mutants. **b**, Developmental stages of *hpl-1; hpl-2* mutants upon RNAi knockdown of transposase encoding ORFs or empty vector control (EV) grown for the same length of time at 23°C. Data from two independent replicates is shown (60-527 worms per treatment in each replicate). Chi-square tests were performed between target RNAi and control (EV) RNAi, and “✓” denotes knockdowns that result in significantly faster growth of *hpl-1; hpl-2* mutants (Bonferroni-adjusted p-value: *p* <0.05).

**Extended Data Figure 8.**
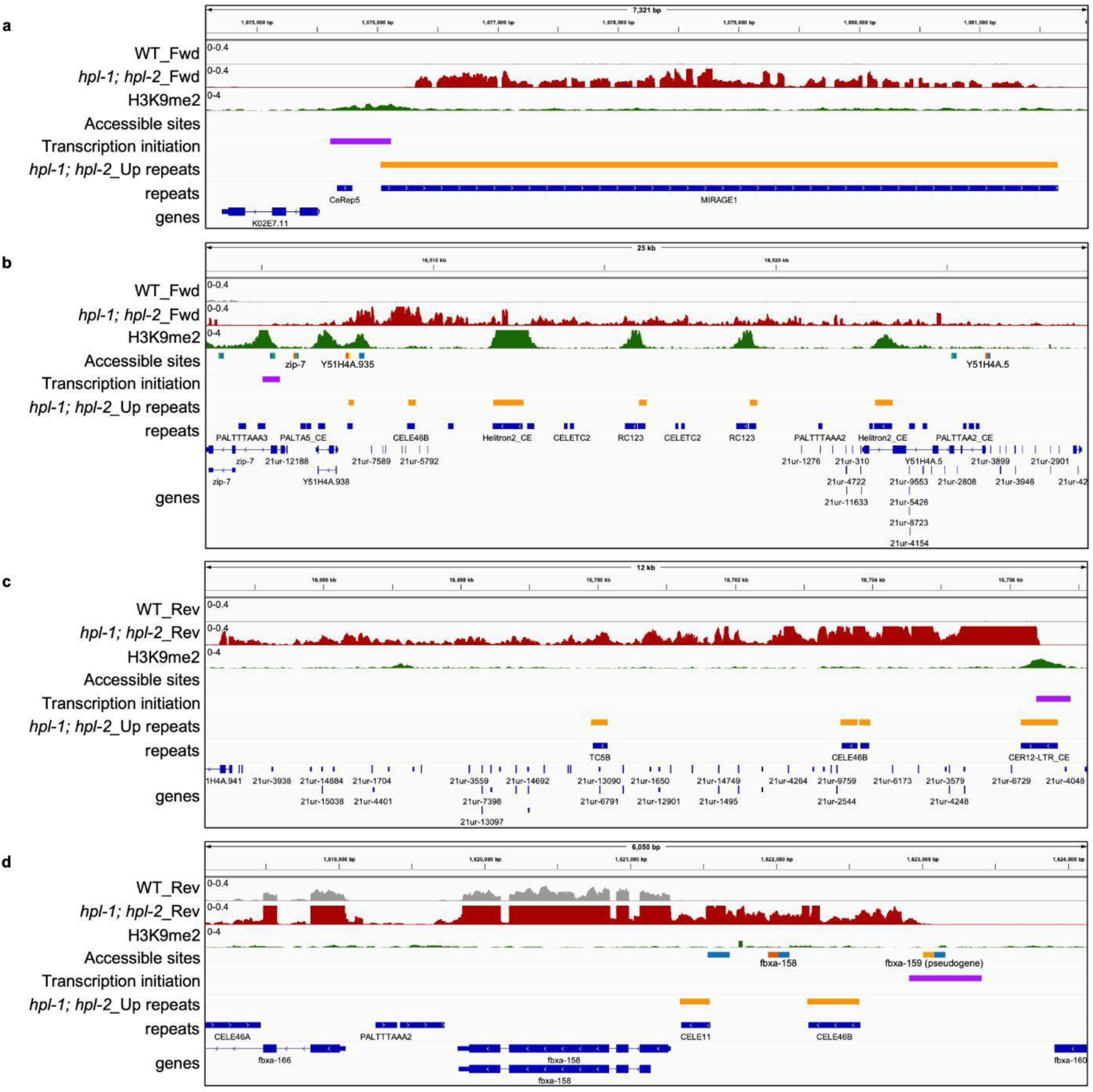
Examples of upregulated repetitive elements in *hpl-1; hpl-2* mutants. **a-d**, IGV browser screenshots of RNA-seq profiles and H3K9me2 ChIP-seq coverage (both in CPM) for representative examples of upregulated repeats in *hpl-1; hpl-2* mutants. **a**, Intrinsically upregulated transposase encoding repeat with H3K9me2 marked transcription initiation site (direct target). **b**, Repeats within a broad upregulated domain, with transcription initiating at an upstream gene promoter marked by H3K9me2 (direct target). **c**, Repeats within a broad upregulated domain, with transcription initiating at an upstream repeat marked by H3K9me2 (direct target). **d**, Repeats within a broad upregulated domain, with transcription initiating at an upstream gene promoter not marked by H3K9me2 (indirect target).

**Extended Data Figure 9.**
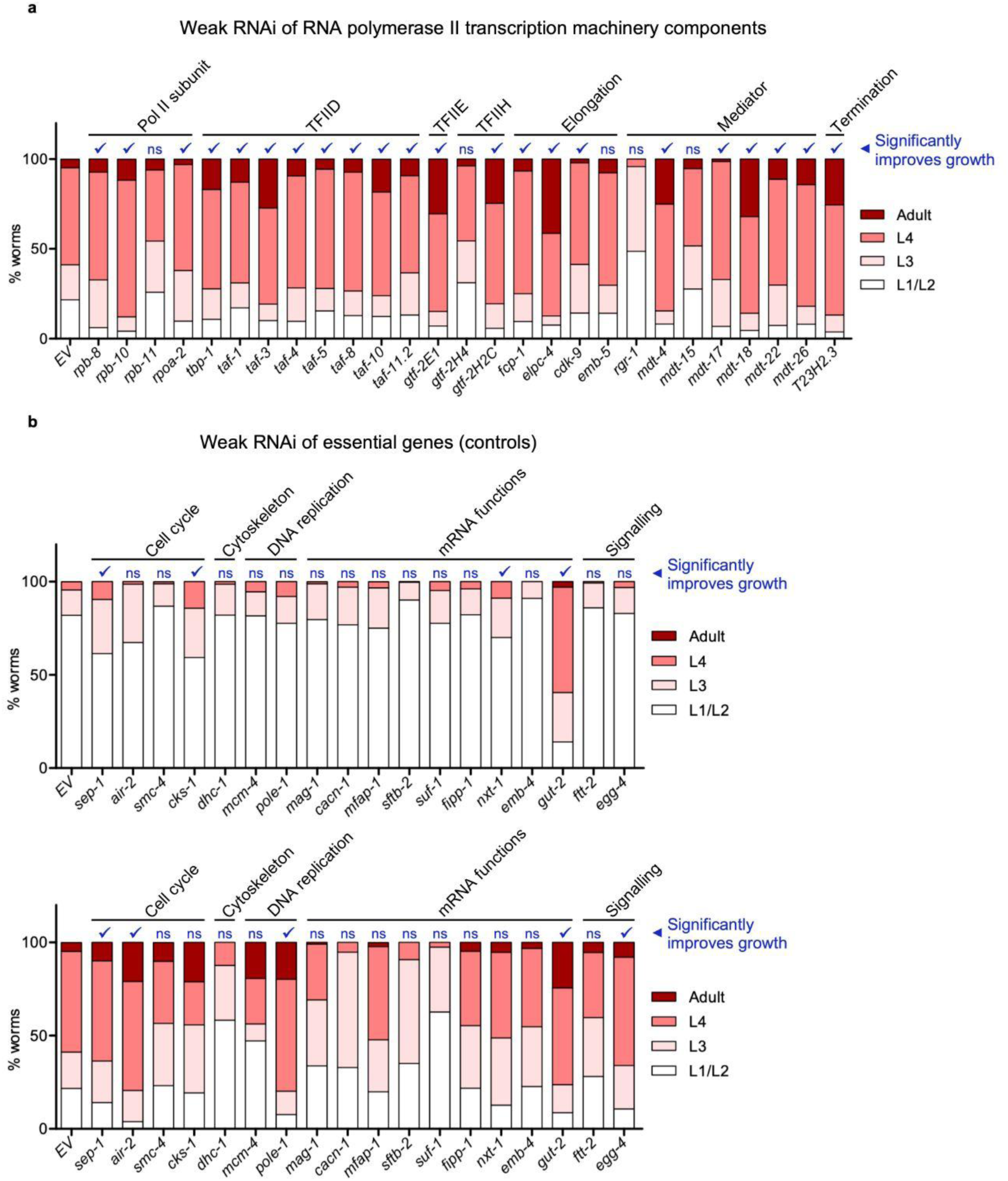
Tests of the effects of weak knockdown of RNA polymerase II machinery and diverse essential genes on *hpl-1; hpl-2* growth rate. **a**, Developmental stages of *hpl-1; hpl-2* mutants upon weak knockdown of various components of RNA polymerase II machinery by RNAi feeding at 23°C. Experimental replicate for Fig. 5a (96-665 worms per treatment in each replicate). **b**, Developmental stages of *hpl-1; hpl-2* mutants upon weak knockdown of diverse essential genes by RNAi feeding at 23°C. Top and bottom show data from two independent replicates (52-665 worms per treatment in each replicate). **a**,**b**, Chi-square tests were performed between target RNAi and control (empty vector, EV) RNAi, and “✓” denotes genes whose knockdown results in significantly faster growth of *hpl-1; hpl-2* mutants (Bonferroni-adjusted p-value: *p* <0.05, ns: not significant).

**Extended Data Figure 10.**
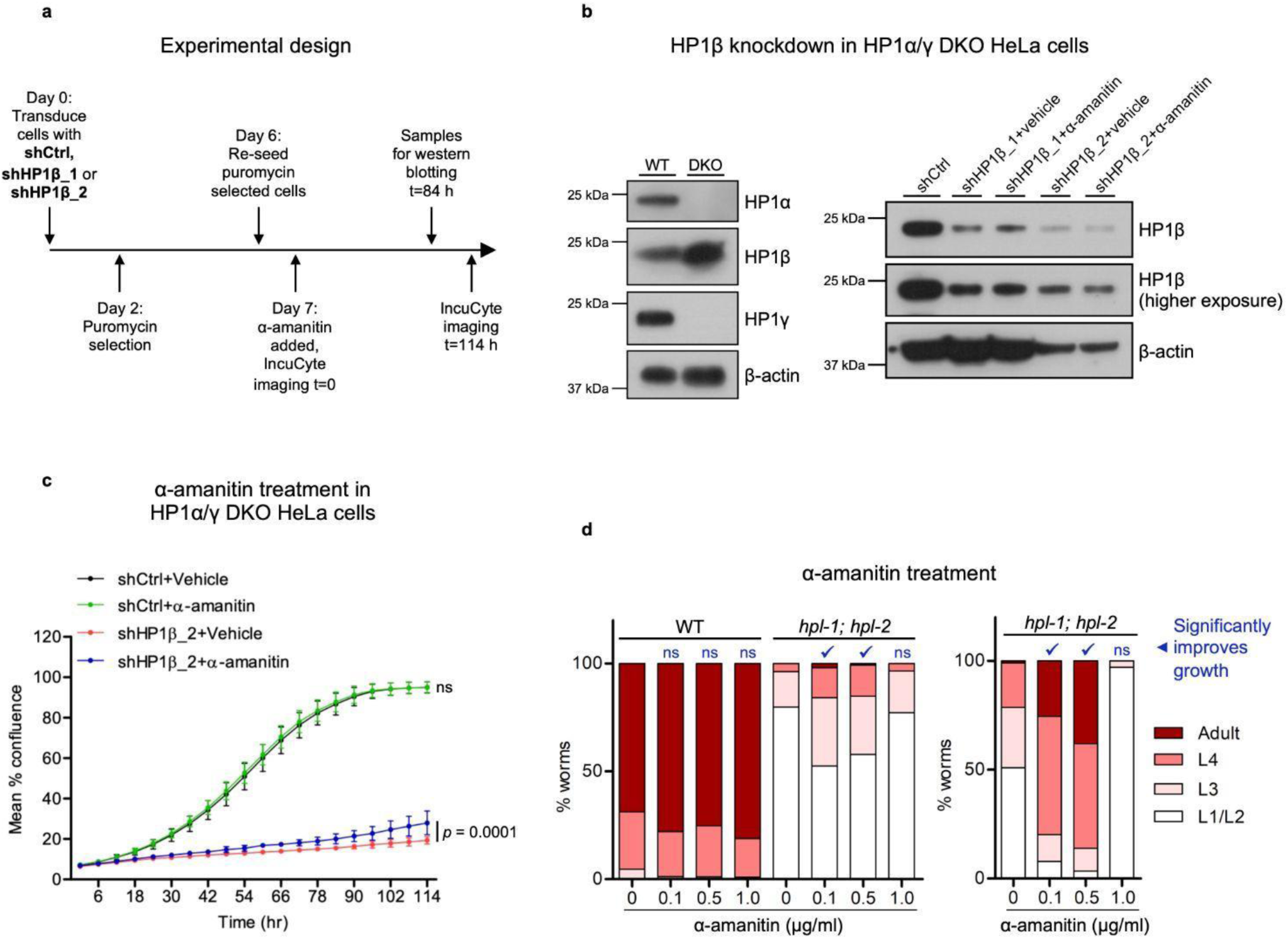
Low-dose α-amanitin treatment improves proliferation of HP1-deficient HeLa cells and growth of *hpl-1; hpl-2* mutants. **a**, Experimental design for testing the ability of low-dose α-amanitin treatment to ameliorate the proliferation defect of HP1-deficient HeLa cells. **b**, Western blot showing absence of HP1α and HP1γ in DKO cells (left) and levels of HP1β knockdown using shHP1β_1 or shHP1β_2 in DKO cells with and without α-amanitin treatment (right). **c**, Mean percent confluence of HP1α/γ double knockout (DKO) HeLa cells with (shHP1β_2) or without (shCtrl) HP1β depletion, treated with 6.25 ng/ml α-amanitin or vehicle (Same experiment as in Fig. 5c, but using an independent shRNA sequence). Each time point shows mean ± standard deviation from duplicate wells. Two-way mixed ANOVA was performed to assess growth trajectory divergence between vehicle and drug treatments: shCtrl+vehicle vs shCtrl+α-amanitin (F = 0.1235, *p* = 1.0) and shHP1β_2+vehicle vs shHP1β_2+α-amanitin (F = 4.086, *p* = 0.0001). Bonferroni post-hoc tests were performed for pairwise differences at each time point: shHP1β_2+vehicle vs shHP1β_2+α-amanitin (*p* <0.05 from 102 hrs onwards). ns: not significant. **d**, Developmental stages of wild-type (WT) or *hpl-1; hpl-2* mutants upon treatment with α-amanitin after growth for the same length of time at 23°C. Experimental replicates for Fig. 5b. Chi-square tests were performed between drug treatments and respective control (0 μg/ml), and “✓” denotes treatments resulting in significantly faster growth (Bonferroni-adjusted p-value: *p* <0.01, ns: not significant).

